# IRF1 Tunes Basal Immunity and Antiviral Readiness in a Context-Dependent Manner

**DOI:** 10.1101/2025.09.16.676475

**Authors:** Eyal Zoler, Irina Miodownik, Shifra Ben-Dor, Daniel Harari, Jiri Zahradnik, Ariel Afek, Gideon Schreiber

## Abstract

Interferon Regulatory Factor 1 (IRF1) plays a pivotal role in interferon (IFN) signaling, yet its context-dependent regulatory functions remain incompletely understood. Here, we dissect the impact of IRF1 on gene regulation in HeLa cells, by targeted knockout (KO) or overexpression (OE) of IRF1. *IRF1* KO did not impair interferon stimulated gene (ISG) expression regulation upon IFN-β stimulation, but partially diminished IFN-γ induced gene regulation. *IRF1* KO did show a homeostatic role in basal gene abundance, including increasing the abundance of some antiviral genes. RNA-seq analysis showed altered expression of both ISGs and immune signaling genes, implicating IRF1 as a dual regulator that fine-tunes gene abundance through both activation and repression. IRF1 OE induced potent antiviral protection in the absence of exogenous IFN, mediated by type I IFN secretion, particularly of IFN-α subtypes. This paracrine effect was confirmed by transcriptomics, cytokine profiling, and mass spectrometry, and was functional even in JAK1-deficient or Ruxolitinib-treated cells but not type I IFN receptor KO cells, suggesting the involvement of non-canonical signaling pathways. Hierarchical clustering of RNA-seq data revealed distinct IFN-independent gene clusters activated or repressed by IRF1, including pathways related to adaptive immunity and T cell function. Using protein-binding microarrays and predictive modeling, we mapped IRF1 binding across promoters and validated functional motifs in the IFIT2 gene promoter by a reporter assay. Our integrative approach establishes IRF1 as a central regulator of antiviral immunity, capable of shaping gene expression both through cytokine signaling and direct promoter binding.

## Introduction

Type I interferons (IFN-I) are essential in initiating both innate and adaptive immune responses against various pathogens (1). They also play a significant role in regulating tumor immunity and contribute to the development of autoimmune diseases. IFN-Is are secreted proteins that induce antiviral activity in almost all nucleated cells in vertebrates, although their antiproliferative and immunomodulatory activities vary with cell type (2–4). The human IFN-I family comprises 17 members, including 13 subtypes of IFN-α, which exhibit high sequence and structural similarity (5–7), along with IFN-β, IFN-κ, IFN-ω, and IFN-ε (8, 9). These IFN-I members all utilize a common receptor composed of the subunits IFN-α receptor 1 (IFNAR1) and IFN-α receptor 2 (IFNAR2) (3, 10).

Upon the engagement of IFN-I with its receptor subunits, a ternary complex is formed, triggering the JAK-STAT signaling pathway. This activation results in the phosphorylation of JAKs and key tyrosine residues on the STAT proteins (11, 12), facilitating their dissociation from the receptor, dimerization, and nuclear translocation where, together with the Interferon Regulatory Factor (IRF)9 they function as transcription factors (13, 14). This process significantly alters the expression of thousands of interferon-stimulated genes (ISGs) (15–17), including a marked upregulation of certain IRFs, especially IRF1 (18).

Interferon-gamma (IFN-γ), which signals through IFNGR is a crucial cytokine primarily produced by natural killer (NK) cells, activated T cells and some antigen-presenting cell subsets is an important component of innate and adaptive immune responses (19–21). It plays a vital role in immune surveillance by enhancing the antimicrobial functions of macrophages and stimulating antigen presentation. The signaling cascade initiated by IFN-γ binding to its receptor activates the JAK-STAT pathway, culminating in the expression of ISGs, many of which mediate antiviral and antibacterial immunity (22, 23). Among the key transcription factors downstream of IFN-γ signaling is IRF1 (24). IRF1 is induced directly by IFN-γ and serves as a master regulator in the immune response. It enhances the expression of various immune-related genes, including those involved in inflammation, apoptosis, and antigen processing(25–27).

Discovered in 1988 by Taniguchi, IRF1 was the first identified member of the IRF family, noted for its transcriptional activation in nuclear extracts post-viral infection (28). IRF1 is a versatile transcriptional regulator critical for various cellular responses, including the host response to viral and bacterial infections, and its involvement in the expression of numerous genes in hematopoiesis, inflammation, immune responses, and cell cycle control (29–31). It acts as both a transcriptional activator and repressor (32–36), binding to interferon-stimulated response elements (ISREs) in the promoters of its target genes (29, 34, 36–38), thus playing an essential role in IFN signaling, immune regulation, tumor suppression, and responses to genotoxic stress (30, 32).

IRFs play critical roles in orchestrating immune responses to viral infections, with IRF9 and IRF1 being central regulators in the type I and type II interferon pathways, respectively. IRF9 is primarily involved in the IFN-I response. It forms a complex with STAT1 and STAT2, known as ISGF3 (interferon-stimulated gene factor 3), which binds to ISREs on target genes, initiating the transcription of a wide array of ISGs(39–42). Although IRF9 and IRF1 are associated with distinct interferon pathways, growing evidence highlights their functional overlap and synergy in modulating antiviral immunity and inflammation(43). IRF1 and IRF9 share the ability to bind ISREs, which enables them to regulate overlapping sets of genes(44). While IRF1 can act independently of IRF9, particularly in IFN-γ-driven immune responses, recent studies show that these two transcription factors often act together to amplify the transcriptional activation of key ISGs(43).

There is compelling evidence that IRF1 suppresses the replication of a variety of RNA viruses and plays a critical role in host antiviral defense, although this can vary depending on the cell type and specific virus. For instance, IRF1 is crucial for activating the transcription of type III IFNs in human hepatocytes infected with Sendai virus(45). However, Type III IFNs, due to their localized receptor abundance and insufficient STAT1 activation, fail to induce IRF1 expression or activate its proinflammatory gene program in epithelial cells. In contrast, the antiviral effects of Type I and II IFNs are more robust(46). In addition to its established antiviral role, IRF1 has been implicated in T cell development, linking it to adaptive immunity and cancer immunosurveillance (47–49). These emerging roles expand the relevance of IRF1 beyond acute infection to tumor immunity and chronic inflammation. Despite this breadth of function, key aspects of IRF1 biology remain unclear, particularly its role in maintaining basal ISG expression and its capacity to function independently of the canonical JAK-STAT pathway. While some studies suggest IRF1 can induce transcription in a STAT-independent manner, the extent and mechanism of this activity are still not sufficiently understood (50).

Similar to other members of the IRF family, IRF1 features an N-terminal DNA-binding domain (DBD) characterized by a sequence of five meticulously conserved tryptophan-rich repeats (51, 52). IRF1 is a regulatory element that plays a central role in the cellular response to IFN signaling. The IRF1 DBD shows a high affinity for the consensus DNA sequence 5’-GAAANNGAA-3’, where “N” represents any nucleotide. Additionally, it can bind to various non-consensus sequences, albeit with lower affinity (53). The IRF1 DBD comprises 136 amino acids with a molecular mass of 16 kDa. This compact domain adopts a helix-turn-helix (HTH) motif, a structural feature commonly found in DNA-binding proteins (54). The HTH motif in the IRF1 DBD facilitates its interaction with the DNA backbone, enhancing its specificity for binding ISRE. The IRF1 DBD binds to the promoters of a broad spectrum of genes, including interferons and genes involved in antiviral protection, and cell growth regulation (29, 32, 34–36). The ability of the IRF1 DBD to interact with diverse promoters emphasizes its extensive influence on gene expression and its significance in regulating cellular responses to IFN signaling. In summary, the IRF1 DBD capacity to recognize and bind specific DNA sequences enables IRF1 to orchestrate the transcriptional regulation of IFN-inducible genes, highlighting its crucial role in maintaining cellular health. Despite extensive knowledge of IRF1’s structure and DNA recognition motifs, relatively little is known about how IRF1 discriminates between promoters under different signaling contexts, or how its function is shaped by chromatin accessibility, cofactor binding, or promoter affinity. Addressing these questions requires a combination of structural, transcriptomic, and functional analyses.

Here, we explore the complex mechanisms underlying IRF1’s regulation of IFN signaling by examining the effects of IRF1 KO and OE in HeLa cells. Through molecular and functional analyses, this research delves into the intricate interactions between IRF1, IFN signaling, and antiviral immunity, offering insights into potential therapeutic strategies for combating viral infections. Notably, our study focuses on IFN-I and IFN-II, which are known for their strong antiviral activity. While IRF1 is well established as a key transcriptional effector of IFN signaling, its precise regulatory functions in maintaining basal ISG expression, modulating cytokine output, and functioning independently of JAK-STAT pathways remain poorly understood. Elucidating these non-canonical roles of IRF1 is particularly relevant given its emerging importance in immune homeostasis and its potential therapeutic relevance in cancer and infection contexts where IFN signaling may be impaired.

## Results

### Loss of IRF1 Disrupts Interferon Signaling Homeostasis and Promotes Constitutive Antiviral Pathway Activation

We have previously shown that *STAT2* KO results in the loss of induction of ISRE-mediated gene expression, while GAS-mediated induction remained unaffected. Only a dual knockout of *STAT2* and *IRF1* fully inhibited type I IFN signaling (11). To further delineate the specific role of IRF1, we generated an *IRF1* KO HeLa cell line. We assumed that the loss of IRF1 would impair the upregulation of gene expression driven by the GAS promoter activated by STAT1 homodimers, while STAT2 KO impairs type I IFN-induced ISGF3 formation. Western blot (WB) analysis confirmed the absence of IRF1 protein independent of IFN-β treatment (Fig. 1A). *IRF1* KO did not significantly impact gene expression following IFN-β treatment (Fig. S1A), however, levels of gene abundance following IFN-γ treatment were lower for many (but not all) IFN-γ induced genes (Fig. 2A). Interestingly, *IRF1* KO altered gene abundance in non-treated cells, as determined by real-time PCR (Fig. 1B) and RNA-seq (Fig. 2B). RNA-seq suggested higher basal abundance of several IRFs, including *IRF7*, and *IRF9* (Fig. S1B and C). To validate that increased mRNA abundance translate to protein abundance of IRF7 and 9 proteins we performed WB, showing a similar trend (Fig. S1E and F).

**Figure 1.**
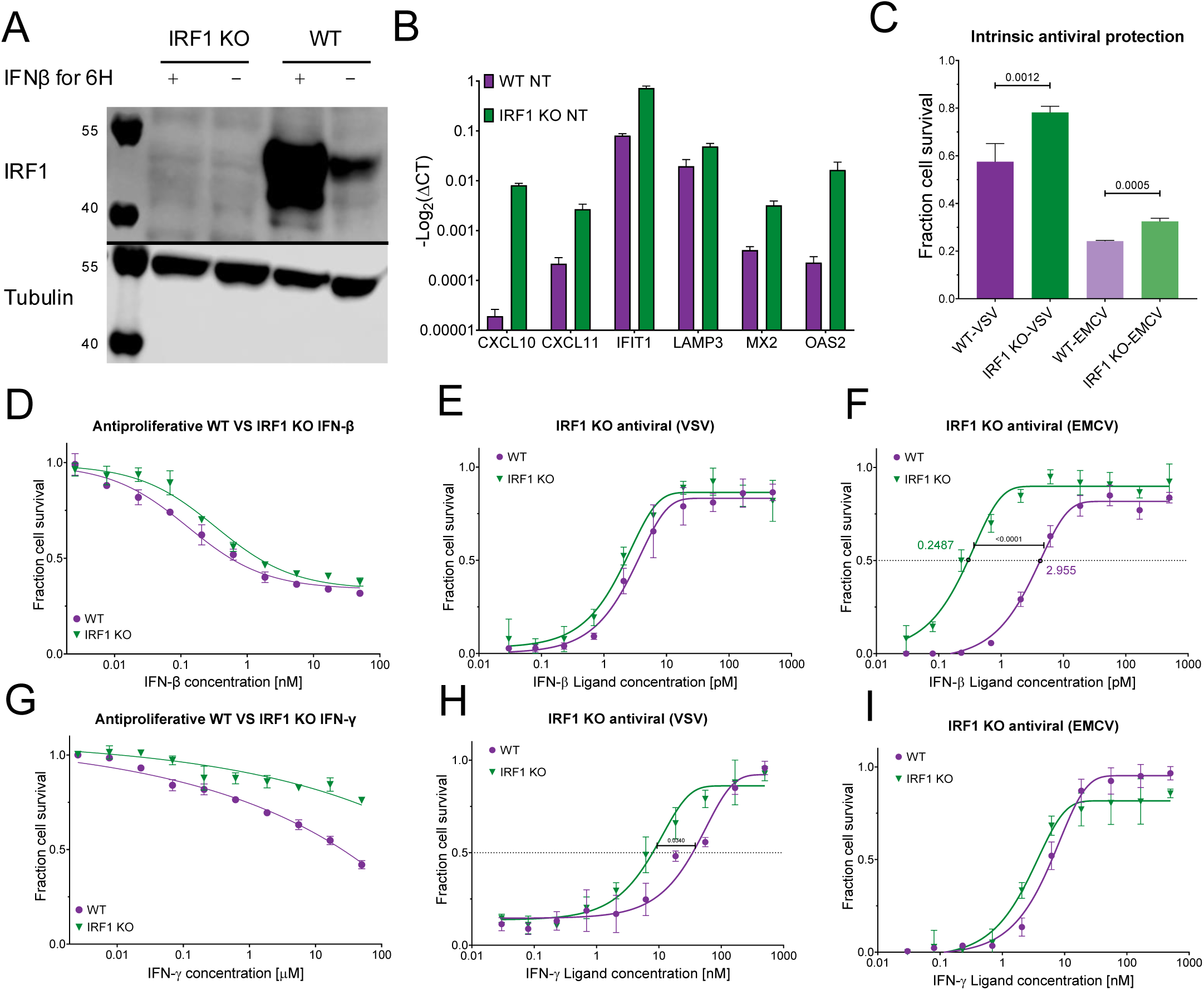
Characterization of *IRF1* KO HeLa cells: protein and gene expression and antiviral responses. (**A**) Western blot analysis of IRF1 protein abundance in WT and IRF1 KO HeLa cells. Quantification of three replicates of this blot is presented in Fig. S1D. (**B**) Relative expression levels of selected ISGs in WT and *IRF1* KO cells, measured by qRT-PCR. Data are shown as – log₂(ΔCT) values normalized to HPRT1. Genes include robustly induced type I IFN targets (XAF1, MX1, MX2, OAS2) and tunable ISGs (IDO1, CXCL10, CXCL11). Data represent mean ± SD of three independent experiments. (**C**) Intrinsic antiviral protection of WT and *IRF1* KO cells following infection with VSV or EMCV in the absence of exogenous IFN. (**D**) Antiproliferative effects of IFN-β. Cells were treated for 96 h and stained with crystal violet to assess viability. (**E-F**) Antiviral activity of WT and *IRF1* KO cells treated with IFN-β for 4 h prior to VSV (18 h) (**E**) or EMCV (20 h) infection **(F)**. **(G-I)** is like **(D-F)** but treated with IFN-γ, for 96 h and stained with crystal violet to assess viability in **(G)** and treated for 8 h in (**H–I**) prior to VSV (18 h) or EMCV (20 h) infection. Cell viability was determined by crystal violet staining. Data represent median of 3–5 independent experiments; error bars indicate SD.

**Figure 2.**
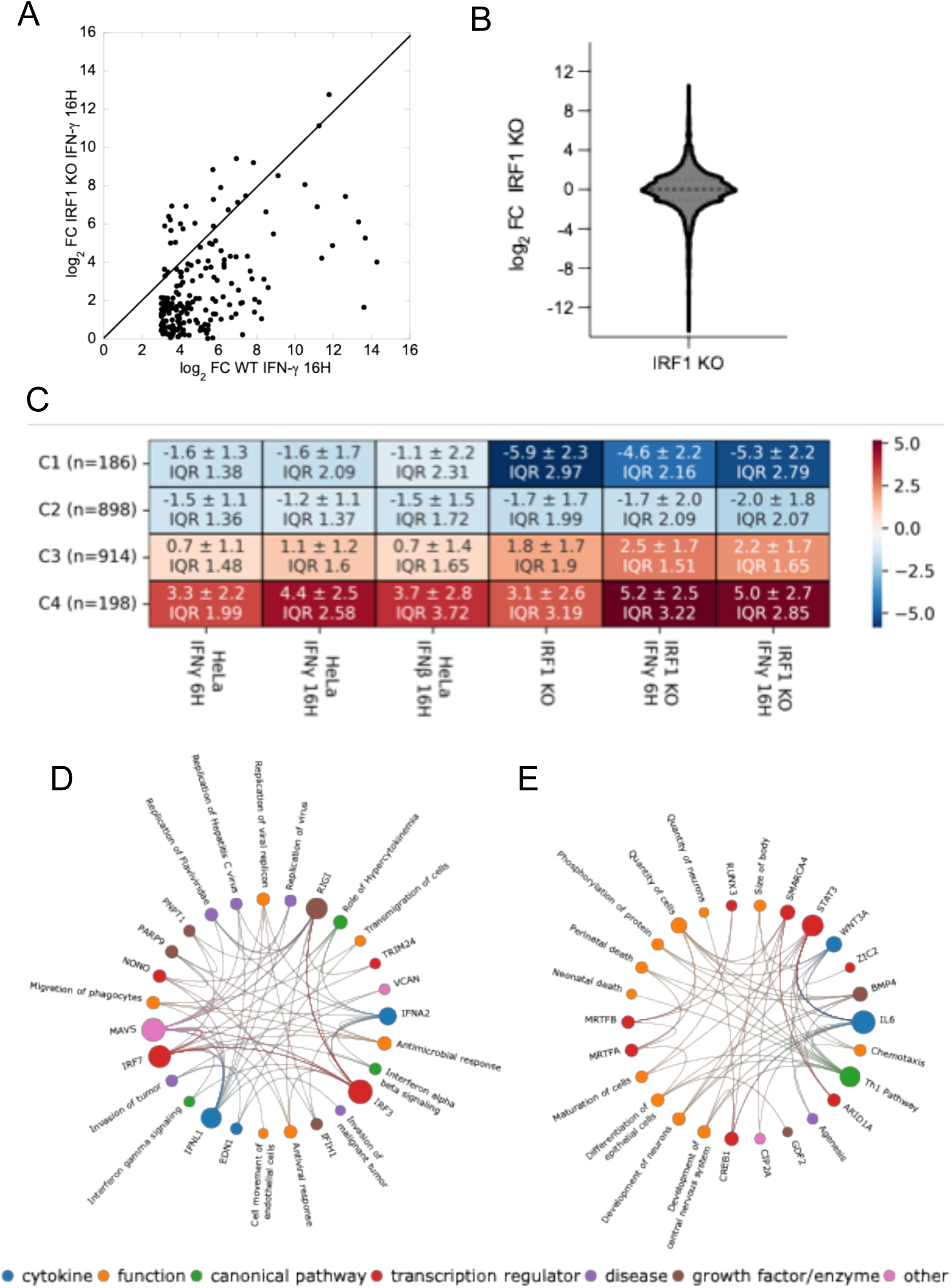
Transcriptomic analysis of *IRF1* KO cells, gene expression patterns and pathway analysis. **(A)** Gene abundance (relative to WT untreated) of *IRF1* KO HeLa cells plotted against WT cells following 16 hours treatment with IFN-γ (100 nM). **(B)** Gene abundance (relative to WT untreated) of *IRF1* KO HeLa cells. **(C)** Heatmap showing hierarchical clustering of genes with significant expression changes (|log2(FC)| > 2) relative to non-treated WT HeLa cells across the following conditions: WT cells treated with IFN-γ (100 nM) for 6 hours, WT cells treated with IFN-γ (100 nM) for 16 hours, WT cells treated with IFN-β (2 nM) for 16 hours, *IRF1* KO cells, *IRF1* KO cells treated with IFN-γ (100 nM) for 6 hours, and *IRF1* KO cells treated with IFN-γ (100 nM) for 16 hours. Clusters are divided into four groups based on median expression, with standard deviation and interquartile range (IQR) calculated for each group. **(D-E)** Pathway analysis of Cluster 4 **(D)** and Cluster 1 **(E)**, performed using QIAGEN Ingenuity Pathway Analysis (IPA). Pathways were selected based on a z-score > 2 and a p-value of overlap < 0.05, indicating significant pathway enrichment in genes upregulated in response to *IRF1* KO and IFN treatments.

Given the substantial alterations observed in basal gene abundance, we performed an antiviral assay using a short viral incubation time without IFN treatment (see Methods, Intrinsic Antiviral Protection). Under these conditions, *IRF1* KO cells exhibited enhanced resistance to both vesicular stomatitis virus (VSV) and encephalomyocarditis virus (EMCV) (Fig. 1C), likely attributable to the elevated basal abundance of key ISGs. Next, we monitored the effect of *IRF1* KO on IFN-β and IFN-γ induced antiproliferative and antiviral potency, showing no effect for the former and only a modest change in the antiproliferative activity upon IFN-γ treatment (Fig. 1D and G). Following, we assessed the antiviral activity of *IRF1* KO cells against VSV and EMCV, after IFN-β and IFN-γ treatments (Fig. 1E, F, H, I). The differences between the WT and KO cells were subtle; however, we observed a slight decrease in the EC50 values in the KO cells, which was statistically significant only for EMCV treated with IFN-β (Fig. 1F). Our hypothesis for the subtle differences between the *IRF1* KO and wild type cells is that the KO exhibits compensatory increased abundance of other IRFs, as shown in Fig. S1E and F.

To gain detailed insight into the transcriptomic changes, we further analyzed our RNA-seq data, comparing WT and *IRF1* KO cells, untreated (NT) or treated with IFN-γ for 6 hours and 16 hours, or IFN-β for 16 hours. WT NT cells served as the baseline for fold-change (FC) analysis. Using the UTAP pipeline (55) and the DESeq2 package (56), we identified significant transcriptomic alterations in the absence of IRF1 (Fig. 2C–E and Fig. S1B). A volcano plot of untreated *IRF1* KO cells (Fig. S1B) revealed higher abundance of key antiviral genes such as *MX1*, *IFIT3*, and *IL6*, alongside lower abundance of genes like *AQP3* and *LPL* (see also Fig. S2). These findings suggest that IRF1 plays a crucial role in maintaining the basal abundance of IFN-regulated genes. Next, we organized the data into a heat map, arranged by hierarchical clustering, which was divided into four distinct groups (Fig. 2C). This approach helped us visualize the gene abundance changes and categorize them. Among the clusters, two stood out as particularly interesting. Cluster 1 constitutes genes whose abundance is strongly decreased in *IRF1* KO cells, with a median log_2_ fold change of approximately -5. Cluster 4 constitutes genes whose abundance is increased in *IRF1* KO cells both without and after IFN treatment, with a median log_2_ fold change of 3-5. To further elucidate the functional significance of these gene clusters, we performed pathway analysis using QIAGEN Ingenuity Pathway Analysis (IPA) (57) for cluster 1 (Fig. 2E), and cluster 4 (Fig. 2D). As anticipated from the IFN-treated WT cells, cluster 4 was strongly enriched for pathways related to MAVS, type I and type II IFN signaling, and antiviral responses. Strikingly, the presence of these same signatures in untreated IRF1 KO cells suggests IRF1 to be a negative regulator of these pathways, suggesting that under steady-state conditions IRF1 also functions as a repressor of MAV activation. Indeed, in Fig. S1C and E we show that IRF7 and IRF9 abundance is increased upon IRF1 KO, which may contribute towards the observed increased expression of other ISGs. In contrast, analysis of cluster 1, which constitutes genes with reduced abundance in *IRF1* KO cells, with or without IFN-γ treatment, revealed strong associations with pathways involved in the IL-6 response and STAT3 signaling, both of which are central to immune regulation. Additionally, we identified in cluster 1 enrichment of the Th1 pathway related genes, which signal through IFN-γ and play a pivotal role in adaptive immunity (58, 59). These findings further underscore the multifaceted regulatory role of IRF1 in orchestrating both innate and adaptive immune responses.

Overall, our results highlight the critical function of IRF1 in fine-tuning IFN signaling pathways and maintaining immune homeostasis. They also offer a mechanistic explanation for the constitutive nuclear localization of IRF1, where it continuously associates with gene promoter elements, even in the absence of specific external stimuli (60).Through this ongoing engagement, IRF1 safeguards the balance of gene expression necessary for preserving cellular integrity and preventing aberrant activation of immune pathways.

### IRF1 Overexpression Confers Antiviral Protection through IFNAR-Dependent but also JAK-Independent pathways

To gain further insight into the role of IRF1 in regulating gene expression, we examined the effects of IRF1 OE through transient transfection. OE of IRF1 resulted in JAK1- and IFNAR-dependent phosphorylation of STAT proteins 48 hours post-transfection (Fig. 3A–C and Fig. S3A), suggesting that IRF1-mediated STAT activation requires an intact type I IFN signaling pathway. The activated STATs in IRF1 OE cells drove antiviral protection against VSV and EMCV even in the absence of IFN-β treatment (Fig. 3D). In contrast, no antiviral effect was detected in IFNAR KO cells, even when IRF1 was overexpressed and IFN-β was added at saturating concentrations (Fig. S3B). Surprisingly, IRF1 OE in JAK1 KO cells resulted in partial antiviral protection (Fig. 3D), despite lack of STAT1 or STAT2 phosphorylation (Fig. 3A), a phenomenon similar to that observed upon MAVS OE, which is known to induce IFN-β secretion (12). Further support for JAK-independent antiviral protection was gained by treatment with the pan-JAK inhibitor Ruxolitinib (61); Even with JAK inhibition, IRF1 OE elicited a partial antiviral response, similar to that observed in JAK1 KO cells (Fig. 3D). To investigate whether antiviral protection was mediated by secreted molecules, we prepared conditioned media (CM) from four different cells: WT, JAK1 KO, IFNAR KO, and GFP OE. Conditioned media were collected 48 hours post-transfection with either IRF1 or GFP (which also serves a control for the effect of transient transfection), centrifuged, and filtered through a 0.22 µm syringe filter to remove cells and cellular debris and to ensure sterility. WT cells were subsequently incubated with the conditioned media, and antiviral activity was assessed. (Fig. 3D). To evaluate the efficacy of the CMs against VSV and EMCV, they were diluted, and antiviral protection was measured (Fig. 3E–F). Except for the GFP CM, which served as a negative control, all other CMs provided full protection. Dilution of the CMs progressively reduced the antiviral activity, confirming that the protective effect was concentration-dependent and mediated by molecules present within the media.

**Figure 3.**
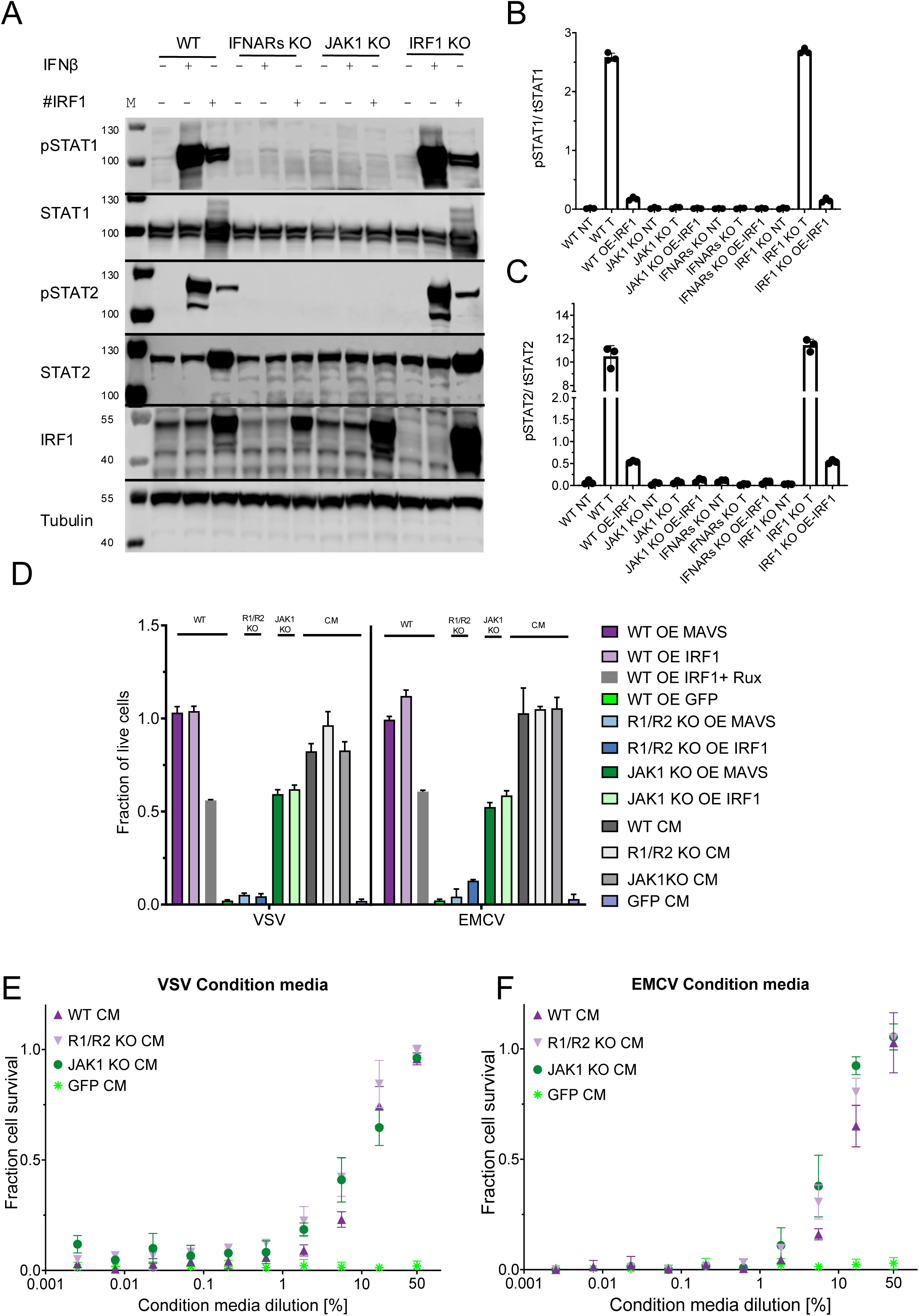
Effects of IRF1 OE on STAT signaling and antiviral activity in various KO cell lines. **(A)** STAT phosphorylation and IRF1 protein abundance in HeLa (WT), *IFNAR* KO, *JAK1* KO and *IRF1* KO cells after 30 min of treatment with 1 nM IFN-β, relative to non-treated cells. #IRF1 designate cells transiently transfected with *IRF1* for 48 hours prior to treatment. **(B)** Normalized pSTAT1 abundance relative to total STAT1 abundance (three replicates). **(C)** Normalized pSTAT2 relative to total STAT2 abundance (three replicates). Quantification of IRF1 is presented in Fig. S3A. **(D)** HeLa WT, IFNAR KO (R1/R2 KO), and JAK1 KO cells were transiently transfected for 48 hours with IRF1 (OE), MAVS (OE), or GFP (OE) (negative control). Where indicated, WT IRF1 OE cells were co-treated with the pan-JAK inhibitor Ruxolitinib (IRF1+Rux). Additional conditions included treatment with conditioned media (CM) collected from WT, *IFNAR* KO (R1/R2 KO CM), *JAK1* KO (JAK1 KO CM), or GFP-transfected (GFP CM) cells. Following transfection or CM treatment, cells were infected with VSV for 18 hours or EMCV for 20 hours. Cell viability was assessed by crystal violet staining and expressed as the fraction of live cells. Bars represent mean ± SD from 3 independent experiments. Statistical significance was determined using one-way ANOVA followed by Tukey’s multiple comparisons test**. (E)** Antiviral activity against VSV for WT HeLa. Cells were treated with CM from WT, *IFNAR* KO, and *JAK1* KO cells for 4 hours before infection with the VSV for 18 hrs. Cells were stained with crystal violet for cell viability. **(F)** Antiviral activity against EMCV for WT HeLa. Cells were treated with CM from WT, *IFNAR* KO, or *JAK1* KO for 4 hours before infection with the EMCV for 20 hours. Cells were stained with crystal violet for cell viability.

These results indicate that IRF1 overexpression induces a robust antiviral response that mirrors the effects of type I IFNs, relying on intact type I IFN receptors in the recipient cells but not in the producer cells. Notably, partial antiviral activity was still observed in the absence of JAK1 both in JAK1 KO cells and in Ruxolitinib-treated cells suggesting that IRF1 can elicit a degree of antiviral protection through a JAK1-independent, paracrine mechanism. Obvious candidates here would be one of the non-canonical IFN induced pathways, including MAPK, PI3K/mTOR and others (62).

### IRF1 Overexpression Induces Secretion of Type I IFNs and Potent Antiviral Factors

The requirement of an intact IFNAR receptor suggests that IRF1 OE induces the secretion of type I IFNs into the medium. Previous studies have suggested a connection between IRF1 and the promoter of IFN-β, though the precise mechanism remains unclear (63). Our goal was to determine whether IRF1 promotes cytokine secretion and to identify the specific cytokines involved. To investigate this, we prepared CM as described above and measured STAT protein phosphorylation by WB. CM from WT and *IFNAR* KO cells OE IRF1 were added to WT and *IFNAR* KO cells for 30 minutes (Fig. 4A and Fig. S4B and C). We observed robust STAT phosphorylation of WT cells for CM from WT and *IFANR* KO cells, whereas no phosphorylation was detected upon adding CM to IFNAR KO cells, confirming the dependence on functional IFNAR signaling. In addition to analyzing STAT phosphorylation, we also examined the effect of CM on IRF1 protein levels. After a 6-hour induction with CM, we detected a marked increase in IRF1 protein levels in WT cells (Fig. 4B and Fig. S4A). To identify which cytokines were present, we utilized the LEGENDplex™ Human Type 1/2/3 IFN Panel (64), which allows for the simultaneous quantification of multiple cytokines, including IFN-α, IFN-β, IFN-λ1, IFN-λ2/3, and IFN-γ (Fig. 4D-E and Fig. S4D-E). In addition to IRF1, we tested MAVS overexpression, known to induce secretion of IFN-β and IFN-λ1 (65, 66). Our findings show high concentrations of IFN-β and IFN-λ1 in cells overexpressing MAVS, with lower concentrations being secreted from KO cells, suggesting a positive feedback loop. MAVS OE resulted in IFN-λ2/3 secretion only from WT cells. IRF1 OE mainly drove IFNα2 secretion, which was higher in *IFNAR* KO or *JAK1* KO cells, suggesting a negative feedback mechanism. A small amount of IFN-β was detected only in JAK1 KO IRF1 OE cells. Next, we evaluated gene abundance of the different type I IFN genes upon IRF1 OE (Fig. 4C) showing that the expression of many of the IFN-α subtypes were increased. Notably, we observed higher expression in the *JAK1* KO IRF1 OE and *IFNAR* KO IRF1 OE cells, providing further evidence of negative feedback of IRF1 activity through type I IFN signaling. Aiming to analyze the CM further, we prepared CM from both WT and *IFNAR* KO cells and after concentration loaded it onto a SEC column (Superdex® 200 10/300 GL), collecting 1 ml fractions. Each fraction was tested for antiviral activity against VSV (Fig. 4F), identifying two active fractions at approximately 37.5 kDa and 24.8 kDa. Mass spectrometry analysis of these fractions revealed three unique peptides. Two peptides from IFN-α1: HDFGFPQEEFDGNQFQK and VGETPLMNADSILAVK (Fig. S5A and S6A) and one peptide from IFN-α4: DRHDFGFPEEEFDGHQFQK (Fig. S5B and S6B). The sequence similarity between IFN-α genes is complicating the analysis required to characterize these cytokines accurately. Together, these findings demonstrate that IRF1 overexpression induces the secretion of type I IFNs, particularly IFN-α subtypes, establishing a paracrine antiviral defense mechanism that amplifies the immune response.

**Figure 4.**
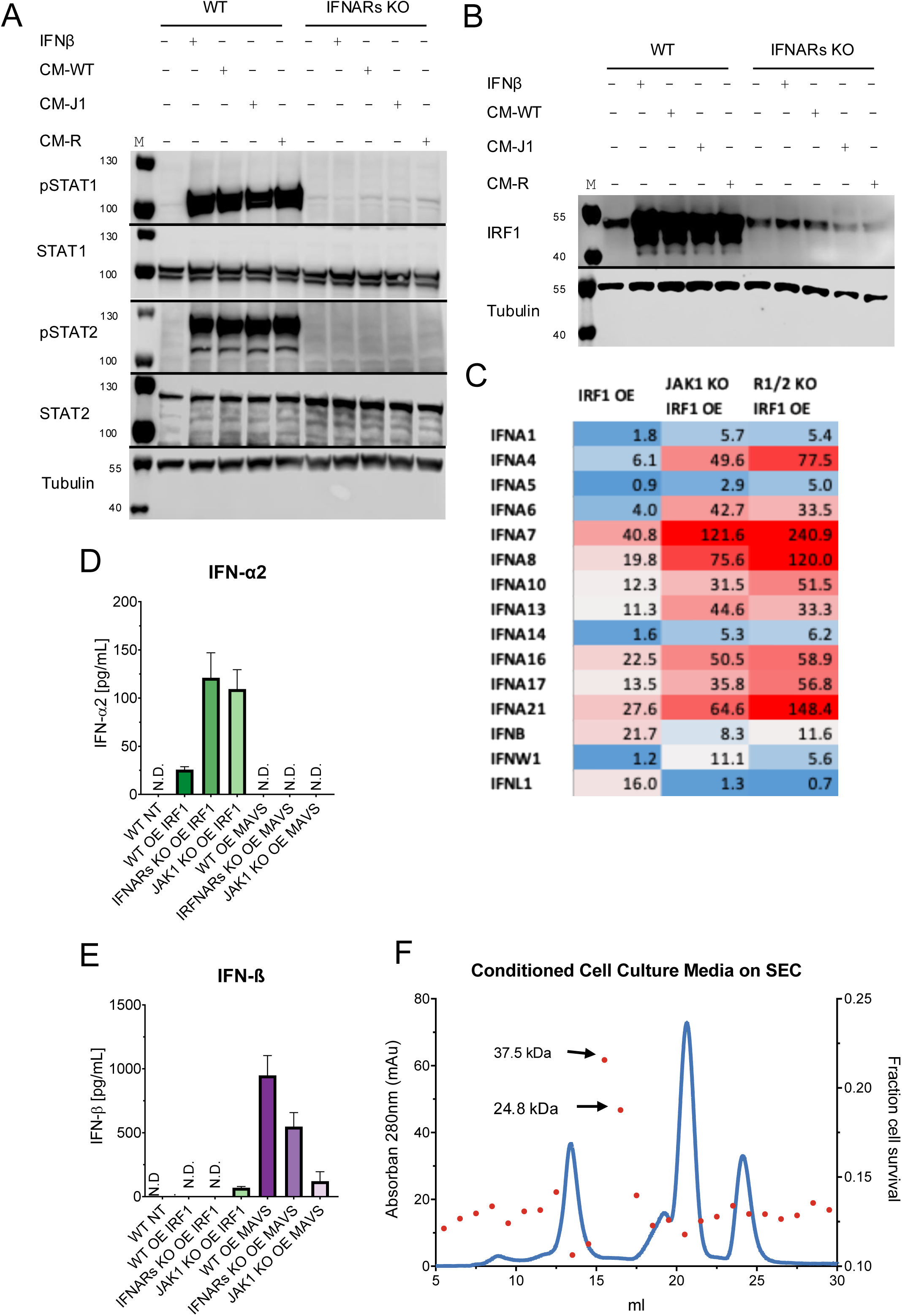
Cytokine profiling and antiviral activity of Conditioned Media from IRF1 OE Cells. **(A)** STAT phosphorylation of HeLa cells (WT) and HeLa *IFNAR* KO after 30 min of treatment with CM from WT, J*AK1* KO (CM-J1) and *IFNAR* KO (CM-R) IRF1 OE cells, relative to non-treated cells. Quantification of this blot and two more replicates is presented in Fig. S4B and Fig. S4C. **(B)** IRF1 abundance in HeLa (WT) and HeLa *IFNAR* KO cells following 6 hours of treatment with CM from WT, *JAK1* KO, and *IFNAR* KO cells, relative to non-treated cells. Quantification of this blot is presented in Fig. S4A. **(C)** Heatmap of the normalized counts of various type I IFNs retrieved from the RNA-seq data. **(D-E)** Flow cytometry analysis of IFN-α2 **(D)** and IFN-β **(E)** abundance in cells overexpressing IRF1 (IRF1 OE) or MAVS (MAVS OE) in WT, *JAK1* KO, and *IFNAR* KO cells. WT NT cells were used as a negative control. **(F)** CM fractionation and their activity. The blue line represents absorption at 280 nm (mAu), while the red dots represent cell survival after treatment with the given fraction of CM for 4 hours, followed by VSV infection for 18 hours. Cell viability was determined by crystal violet staining.

### IRF1 Orchestrates Both Interferon-Dependent and Independent Immune Gene Programs

To further dissect the molecular consequences of IRF1 overexpression and to identify the broader gene expression changes involved, we transfected IRF1 for 48 and extracted RNA from WT IRF1 OE, JAK1 KO IRF1 OE, and IFNAR KO IRF1 OE cells for RNA-seq analysis. Plotting log₂(FC)| > 2.5 of IFN-β (Fig. S7A) or IFN-γ treated cells (Fig. S7B) in relation to IRF1 OE cells shows that indeed gene abundance upon IFN-β treatment mimics that observed upon IRF1 OE. For a more detailed analysis, untreated WT, JAK1 KO, and IFNAR KO cells served as baseline controls for FC calculations. Differential expression analysis was performed using the UTAP pipeline (55) and DESeq2 package(56). Additionally, for comparison, we included WT cells treated with 1 nM IFN-β for 16 hours to assess IFN-induced gene activation, and WT cells overexpressing GFP (WT GFP OE) as a negative control to account for any transfection-induced upregulation of type I IFN responses (57). All genes exhibiting significant expression changes relative to baseline (|log₂(FC)| > 2) were extracted and organized into a heatmap. The heatmap was generated by hierarchical clustering and divided into four distinct groups based on expression patterns. For each cluster, the median expression and interquartile range (IQR) were calculated (Fig. 5A). The most notable expression patterns were observed in clusters 3 and 4, which exhibited distinct regulatory characteristics. Cluster 4 shows strong upregulation in both WT IRF1 OE cells and WT cells treated with IFN-β, consistent with interferon-mediated gene induction. This upregulation was absent in JAK1 KO and IFNAR KO cells and remained comparable to the GFP control, supporting the interpretation that cluster 4 genes represent classical interferon-stimulated genes dependent on intact type I IFN signaling. In contrast, cluster 3 comprises of genes that were robustly upregulated in IRF1 OE cells irrespective of whether JAK1 or IFNAR are present, indicating that their expression is independent of type I IFN signaling (for specific genes of cluster 3 see Fig. S7C). These findings suggest that IRF1 can initiate a distinct transcriptional program that bypasses the canonical IFN –JAK–STAT pathway. Pathway analysis of cluster 3 was performed using QIAGEN Ingenuity Pathway Analysis (IPA) (57). This analysis revealed significant enrichment of pathways related to the innate immune system, including genes such as IL1B, IL3, and the STAT6 signaling pathways (Fig. 5B). In addition, strong activation of pathways associated with the adaptive immune system was observed, including “Interruption of T lymphocytes” and “Adhesion of mononuclear leukocytes,” among others. Thus, beyond its role in inducing type I IFN secretion and upregulating ISGs, IRF1 overexpression can also directly activate genes involved in the innate and adaptive immune systems. This dual activity may help explain the partial antiviral protection observed in *JAK1* KO IRF1 OE cells, even in the absence of canonical type I IFN signaling. Another interesting observation emerged from the Venn diagram analysis (Fig. 5C), where we compared the different clusters. We identified 142 distinct genes that were upregulated upon IRF1 OE and 157 genes upon IRF1 KO. However, the abundance of 41 genes were increased both by IRF1 OE and IRF KO, highlighting that IRF1 can act both as a transcriptional activator and a repressor on the same gene.

**Figure 5.**
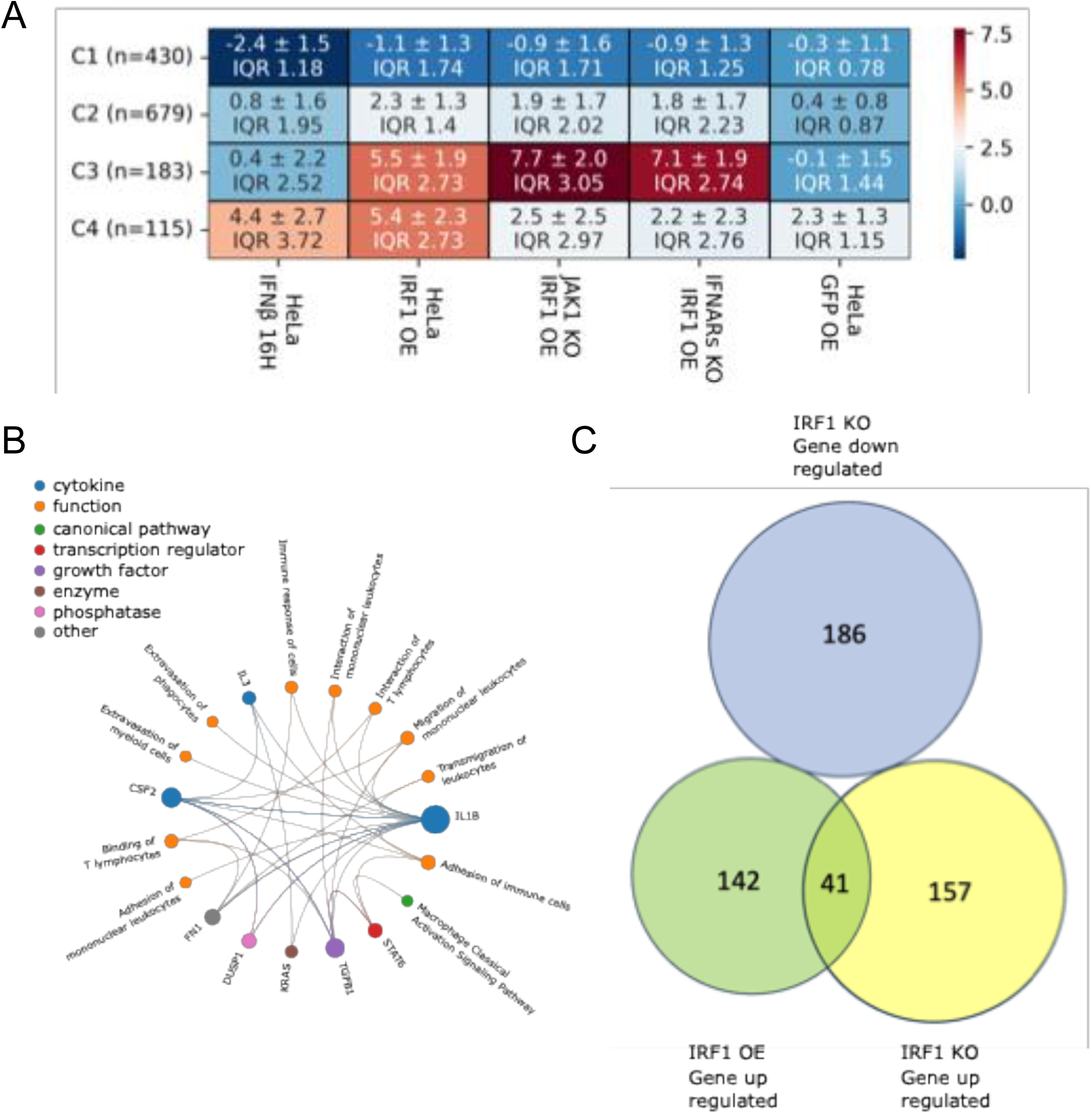
Gene regulation and signaling pathway alterations upon IRF1 overexpression. **(A)** Heatmap showing hierarchical clustering of genes with significant expression changes (|log2(FC)| > 2) across WT IRF1 OE, *JAK1* KO IRF1 OE, and *IFNAR* KO IRF1 OE cells 48 hours post-transfection, with WT cells treated with 2 nM IFN-β for 16 hours and WT GFP OE used as control. Clusters are divided into four groups based on median expression. Standard deviation and IQR were calculated for each group. **(B)** Pathway analysis of Cluster 3 was performed using QIAGEN Ingenuity Pathway Analysis (IPA). Pathways were selected based on a z-score > 2 and a p-value of overlap < 0.05, indicating significant pathway enrichment in genes upregulated by IRF1 OE. **(C)** Venn diagram comparing gene sets from different clusters, showing genes upregulated in *IRF1* KO (Cluster 4 Fig. 2) cells versus those upregulated in IRF1 OE (Cluster 3 Fig. 5) cells and downregulated in *IRF1* KO (Cluster 1 Fig. 2).

Collectively, these results demonstrate that IRF1 modulates immune gene expression through two parallel mechanisms: by driving interferon-dependent antiviral programs and by directly activating innate and adaptive immune pathways independently of canonical type I IFN signaling.

### Characterization of IRF1 DNA-Binding Specificity Using Protein Binding Microarrays

We hypothesized that the distinct regulatory effects of IRF1 on transcription as observed upon its KO or OE may arise from variations in its DNA binding affinity to different promoter regions. To explore this possibility, we aimed to determine IRF1 binding to promoter regions of type I IFN-induced genes and genes differentially regulated by IRF1 OE and KO. At first, we verified the DNA binding motif (DBM) of IRF1 by performing Protein Binding Microarray (PBM) experiments (67, 68). For this, we purified the IRF1 DNA Binding Domain (DBD), comprising amino acids 1 to 136 (MW16 kDa). To enable fluorescence-based detection, the DBD was conjugated to mNeonGreen, a fluorescent protein with a MW of 26 kDa, resulting in a fusion protein of 42 kDa. The fusion protein was expressed in *E. coli* and purified by cation exchange chromatography using SP Sepharose, taking the advantage of the high isoelectric point (pI = 10.2) of the DBD for direct column binding, followed by elution with high-salt buffer (Fig. S8A). The protein was further purified by SEC using a Superdex 75 PG column (Fig. S8B). Following purification, we evaluated the DNA binding specificity of the fusion protein. Two double-stranded DNA oligonucleotides were synthesized: one containing a previously reported IRF1 consensus binding sequence (5’-GAGAAGTGAAAGTACTTTCACTTCTC-3’) (54), and a scrambled control sequence with identical nucleotide composition (5’-ATATTACGTCGCACTAGGATAGATCT-3’). Each oligonucleotide had an approximate MW of 16 kDa. Mass photometry analysis was conducted on three samples: IRF1 DBD-mNeonGreen without DNA, with non-specific (scrambled) DNA, and with specific target DNA (Fig. 6A). The IRF1 fusion protein alone exhibited a mass of approximately 45 kDa. Adding scrambled DNA produced a slight increase to ∼50 kDa (likely reflecting minor non-specific interactions), and adding specific target DNA resulted in a mass of ∼65 kDa. This confirmed the specific and stable binding of IRF1 DBD to its target DNA sequence, despite the highly positive charge of the DBD. Subsequent size measurements were conducted by SEC on a Superdex® 200 10/300 GL column. Three distinct wavelengths were monitored: 280 nm, corresponding primarily to protein absorbance; 260 nm, indicative of nucleic acid (DNA) absorbance; and 498 nm, corresponding to the fluorescence emitted by the mNeonGreen tag conjugated to the IRF1 DBD. The IRF1 DBD-mNeonGreen fusion protein was incubated with the specific DNA sequence for 4 hours prior to SEC analysis (Fig. 6B). Parallel measurements were performed using scrambled DNA and in the absence of DNA as controls (Fig. S9A–C). SEC analysis of the specific DNA-bound sample revealed a major peak corresponding to an apparent molecular mass of ∼58 kDa, with a dominant 260 nm signal relative to 280 nm, indicative of the fusion protein bound to DNA. A second prominent peak was detected at ∼42 kDa, with a higher 280 nm signal, corresponding to unbound IRF1 DBD-mNeonGreen. A third peak at ∼16 kDa, characterized by a dominant 260 nm signal, corresponds to free DNA oligonucleotides. Next, we assessed the thermal stability of the IRF1 fusion protein under different binding conditions: without DNA, with scrambled DNA, and with specific DNA. As shown in Fig. 6C, binding to the specific DNA sequence resulted in a higher melting temperature, consistent with the increased stability typically observed in protein–DNA complexes compared to the unbound protein. Having confirmed the functionality and stability of the IRF1 DBD-mNeonGreen fusion protein, we proceeded with the PBM experiment to characterize the preferences of IRF1-DBM to bind DNA. The PBM analysis successfully identified a sequence-specific binding motif, which was visualized as an energy-normalized sequence logo using the enoLOGOS software (69) (Fig. 6D). These results confirmed the sequence specificity of IRF1 DNA binding and provided a foundation for predictive modeling of IRF1 occupancy across promoter regions of interferon-regulated genes.

**Figure 6.**
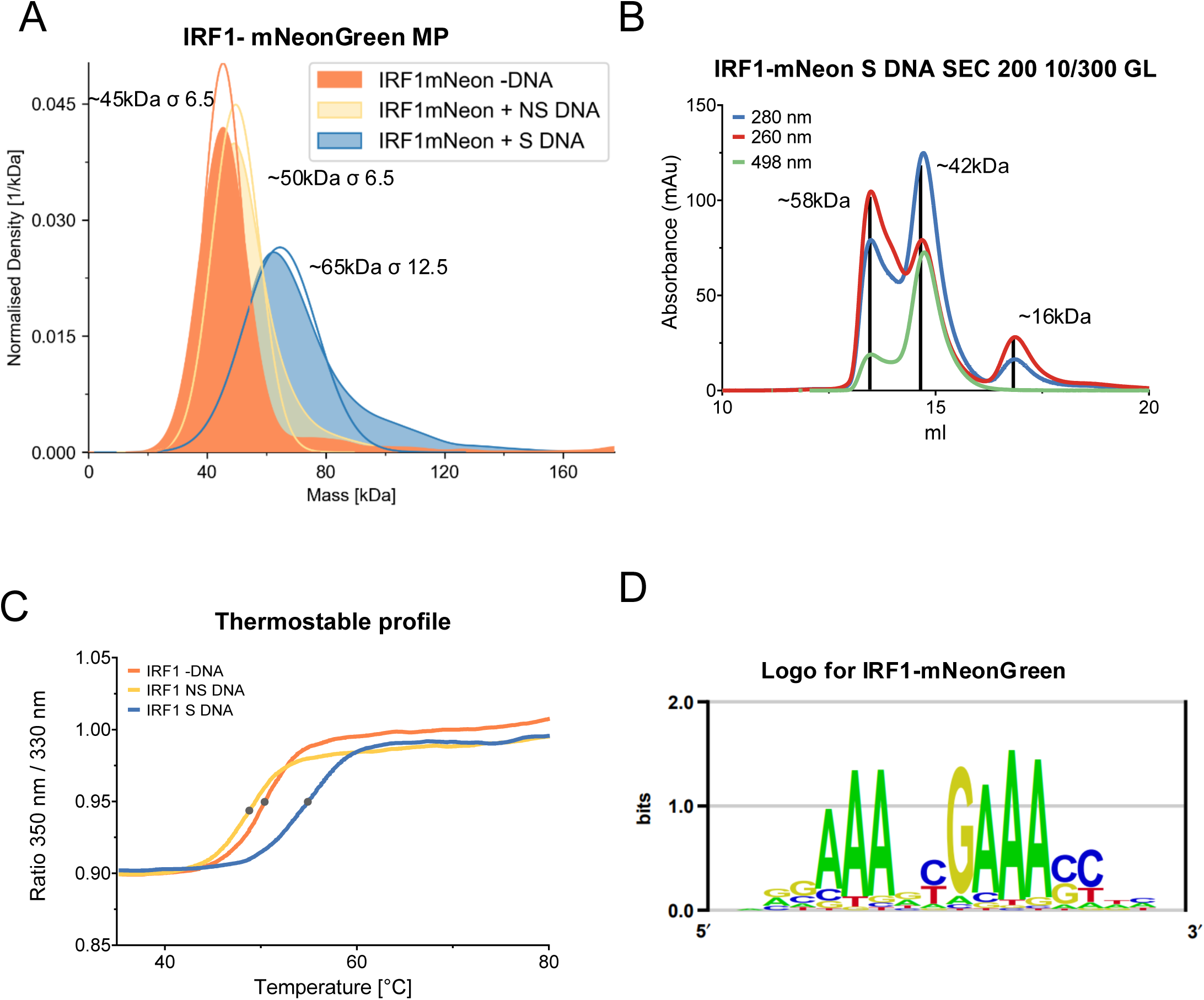
Verification of IRF1 DNA-binding specificity using mass photometry and size exclusion chromatography. **(A)** Mass photometry of the IRF1-mNeonGreen without DNA (-DNA) with scramble DNA (+NS) and with specific DNA (+S DNA). The data shown here represents one example from three independent biological replicates. **(B)** Size exclusion chromatography of IRF1 mNeon incubate for 4 hours with specific. The blue line shows absorbance at 280 nm, the red line 260 nm, and the green line 498nm. **(C)** Thermal Stability of IRF1-mNeonGreen without DNA (-DNA) with scramble DNA (+NS) and with a specific DNA (+S DNA). **(D)** The energy-normalized binding motif logo for IRF1-mNeonGreen was generated using the enoLOGOS software (69).

### Predictive Modeling of IRF1 Promoter Occupancy Reveals Functional Binding Sites in Antiviral Gene Networks

To further investigate IRF1 binding dynamics, we developed a predictive model based on binding affinities derived from DNA sequence data. Using the output of our PBM analysis, we first generated a sequence logo and calculated binding affinities for all possible 7-mer sequences, representing potential IRF1 recognition motifs. We then analyzed the promoter regions of genes identified through our RNA-seq clustering analysis, focusing on three key groups: 1. Genes which abundance was reduced in cluster 1 in *IRF1* KO cells, representing genes were IRF1 is a transcription factor, independent on IFN (Fig. 2A); 2. genes which abundance increased upon *IRF1* KO (cluster 4), representing genes which expression is repressed by IRF1, but induced by IFN. 3. Genes which abundance is increased upon IRF1 OE (cluster 3, Fig. 5A), corresponding to genes were IRF1 is an inducer of transcription, independent on IFN. For each gene of interest, we extracted its promoter region defined as 1,000 base pairs upstream and 500 base pairs downstream of the transcription start site, adjusted for gene orientation. Using the PBM-derived binding affinities, we constructed a gene-specific model predicting IRF1 binding strength and location across each promoter region (Fig. S10). This model allowed us to identify high-confidence candidate binding sites for the IRF1 protein. To validate the model’s predictive power, we re-analyzed IRF1 ChIP-seq data from the Gene Expression Omnibus (GEO) database (GSM6928615, GSM6928616) (70), which originated from a previous study examining time-dependent recruitment of GAS, ISGF3, and IRF1 complexes to GAS, ISRE, and composite elements. Data was processed using the Galaxy platform (71), and visualized with the Integrated Genome Browser (IGB) (72) (Fig. 7 and Fig. S11). Although the original ChIP-seq was performed in Huh7.5 cells, we adapted the analysis to HeLa cells to align with our experimental system. Remarkably, our analysis revealed strong ChIP-seq peaks that overlapped with high-affinity binding sites predicted by our model (Fig. 7A–D, Fig. S11A–D), supporting the model’s accuracy in identifying bona fide IRF1 regulatory elements.

**Figure 7.**
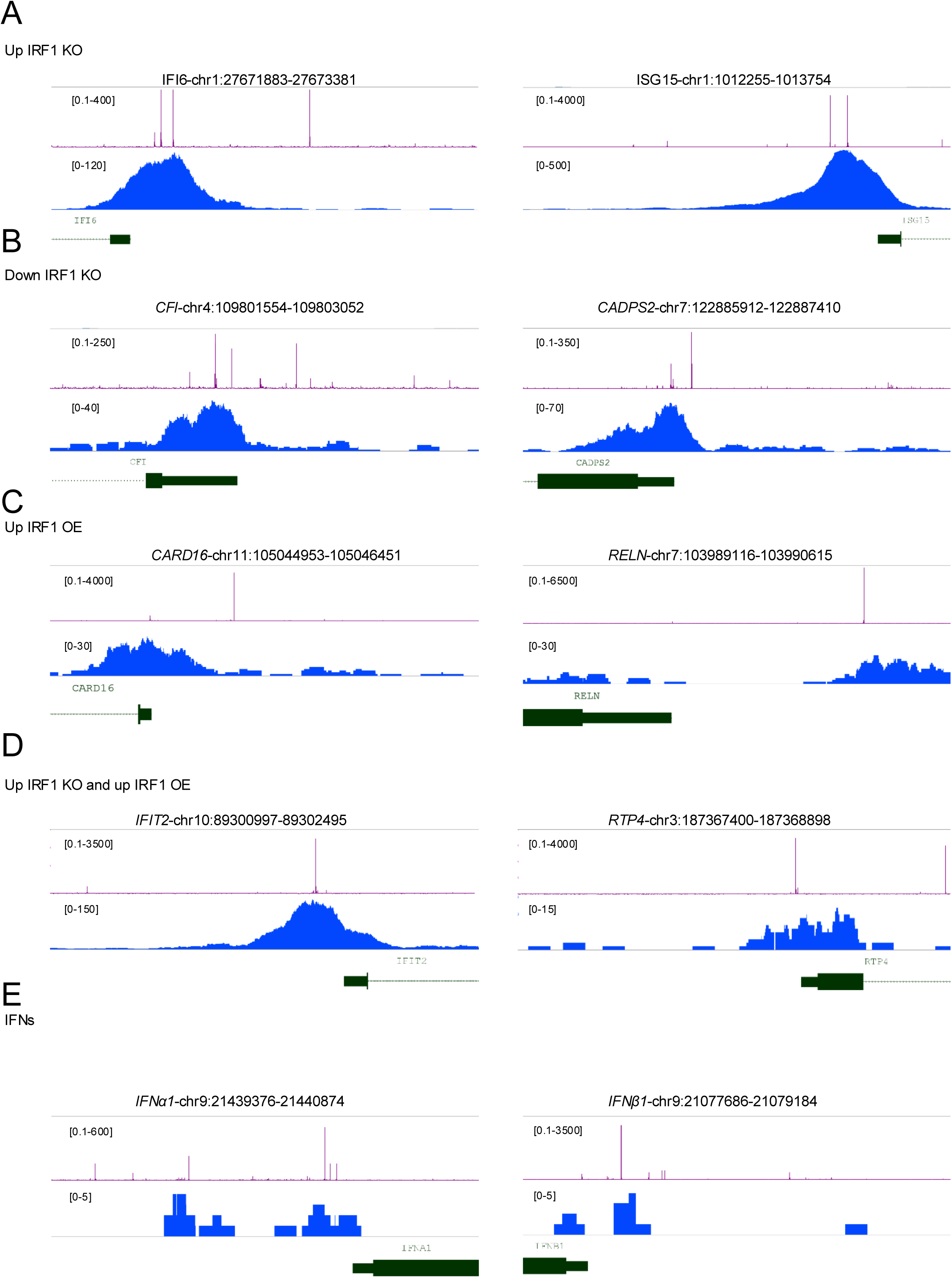
Predicted and observed IRF1 binding in promoter regions of key immune-related genes. **(A-E)** Comparison of predicted IRF1 binding affinity and ChIP-seq coverage in promoter regions of genes identified from RNA-seq analysis. The promoter regions include 1,000 base pairs upstream and 500 base pairs downstream of the transcription start site. Predicted IRF1 binding affinity is shown in purple and was visualized using inverse log₂-transformed z-scores (2^z^-score) to represent the relative strength of predicted interactions on a linear scale. Raw ChIP-seq data were reprocessed with a standardized pipeline from GEO datasets (GSM6928615, GSM6928616) represented in blue, showing IRF1 binding coverage across the promoter regions. Gene annotations are in green. **(A)** Genes showing upregulation in the *IRF1* KO, corresponding to cluster 4 from RNA-seq analysis (Fig. 2C). **(B)** Genes showing downregulation in the *IRF1* KO, corresponding to cluster 1 from RNA-seq analysis (Fig. 2C). **(C)** Genes showing upregulation in IRF1 OE, corresponding to cluster 3 from RNA-seq analysis (Fig 5A). **(D)** Genes showing upregulation in both the *IRF1* KO and IRF1 OE, illustrating genes that are sensitive to changes in IRF1 levels(Fig. 5C). **(E)** Analysis of the IFN promoter region, where a subtle ChIP-seq peak aligns with predicted binding sites. Although this peak is less pronounced, it may be functionally significant due to the high affinity of IFN receptors, which can initiate an antiviral response even with low-level IFN expression. More genes are shown in Fig. S11.

Notably, while the type I IFN promoter regions lacked a dominant ChIP-seq peak in the original analysis, our model predicted high affinity binding sites (Fig. 7E, Fig. S10E, Fig. S11E). Closer inspection revealed a small but distinct ChIP-seq signal aligning with the predicted sites an observation that is biologically meaningful given that even minimal IFN expression can trigger a potent antiviral response due to the picomolar concentrations needed to activate a biological response (73, 74). Thus, weak DNA binding may be sufficient to initiate a biologically relevant response, supporting the value of our model in identifying subtle yet functional regulatory elements.

Collectively, our findings demonstrate that IRF1 binding is highly sequence-specific and that its promoter occupancy can be accurately predicted using a binding affinity-based model. By integrating these predictions with re-analyzed ChIP-seq data, we identified both prominent and subtle IRF1 binding events across key immune gene promoters, including type I IFNs particularly IFN-α subtypes and canonical ISGs. These results establish IRF1 as a central transcriptional regulator of antiviral immunity, capable of directly modulating both IFN-α expression and downstream immune gene networks. This provides a mechanistic framework for understanding the distinct regulatory effects of IRF1 under KO and OE conditions.

### Targeted Promoter Analysis of IFIT2 Reveals Functionally Predictive IRF1 Binding Sites

To experimentally validate our predictive binding model, we selected IFIT2 as a representative target gene for detailed analysis. IFIT2 (Interferon-Induced Protein with Tetratricopeptide Repeats 2) is a well-characterized type I IFN ISG, which expression is rapidly induced following IFN-α/β signaling. Its role in antiviral defense is modulating cellular responses to viral RNA, with its transcription being tightly regulated at the promoter level (75–77). We chose IFIT2 because it exhibited consistent upregulation under multiple conditions: in IRF1 KO cells, in IRF1 OE cells, and following IFN-β treatment. Notably, increased IFIT2 expression was also observed in both JAK1 KO IRF1 OE and IFNARs KO IRF1 OE cells (Fig. 8A, log₂ fold change from RNA-seq), highlighting its robust transcriptional activation across distinct genetic backgrounds and signaling contexts. Using our IRF1 promoter binding model, we analyzed the IFIT2 promoter region (chr10:89300997–89302495), defined as 1,000 base pairs upstream and 500 base pairs downstream of the transcription start site. The model identified two high-affinity IRF1 binding peaks near the start codon, which is marked in green in Fig. 8B. A focused view of this region (chr10:89301927–89301996, ∼70 bp) is shown in Fig. 8C (WT), highlighting the location of the two predicted binding sites.

**Figure 8.**
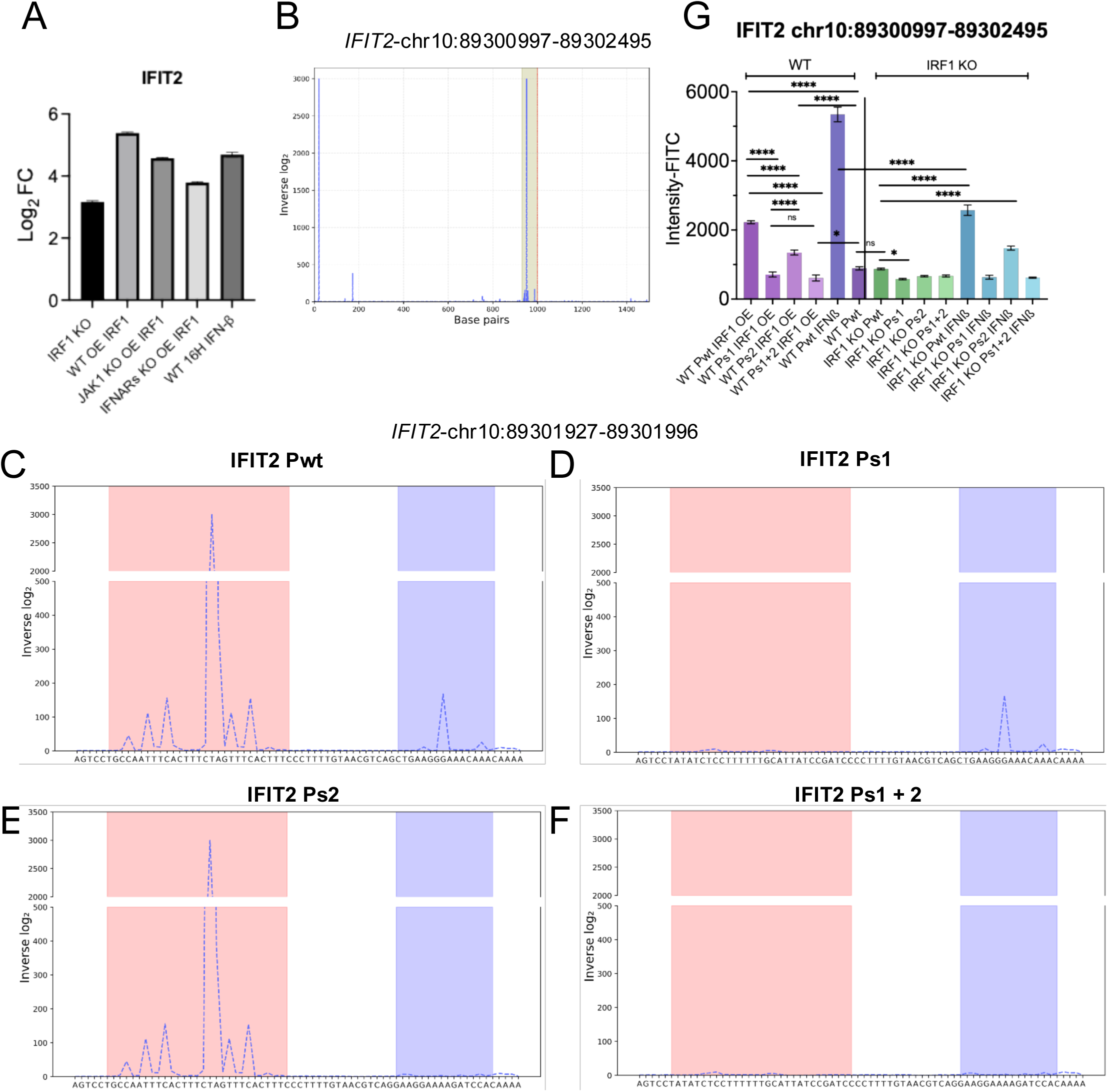
Functional validation of predicted IRF1 binding sites in the IFIT2 promoter. (**A**) Normalized RNA-seq expression of IFIT2 in HeLa cells under the indicated conditions: *IRF1* KO, IRF1 OE, JAK1 KO + IRF1 OE, *IFNAR* KO + IRF1 OE, and WT treated with IFN-β for 16 hours. (**B**) Predicted IRF1 binding affinity across the IFIT2 promoter (chr10:89300997–89302495), based on inverse log₂-transformed z-scores. The analyzed region includes 1,000 bp upstream and 500 bp downstream of the transcription start site. Two predicted binding sites near the start codon are highlighted in green. (**C–F**) Predicted IRF1 binding sites on the WT IFIT2 promoter (**C**), and three scrambled variants in which one or both predicted IRF1 binding sites were scrambled: Ps 1 (**D**), Ps 2 (**E**), and Ps 1+2 (**F**). Binding affinity was recalculated for each sequence and plotted using inverse log₂-transformed z-scores. (**G**) Reporter assay results for the IFIT2 promoter fussed with the eUnaG2 gene expressed in WT and IRF1 KO HeLa cells. Cells were transfected with reporter constructs containing the WT promoter or scrambled variants and were either untreated, transfected with IRF1 OE, or treated with IFN-β for 12 hours. Reporter expression is shown as fold change in fluorescence intensity measured by flow cytometry. Data represent mean ± SD from independent experiments, from which significance was calculated using one way Anova with post Tukey test.

To evaluate the contribution of each predicted site, we generated three scrambled versions of the sequence: one with the first, higher peak scrambled (Ps1, Fig. 8D), one with the second (lower) peak scrambled (Ps2, Fig. 8E), and one with both sites scrambled (Ps 1+2, Fig. 8F). The predictive model was re-run on each of the modified sequences, alongside the wild-type control. As shown in Fig. 8C–F, scrambling of either site reduced the predicted IRF1 binding potential, and scrambling both eliminated it entirely. Notably, the first peak exhibited a higher predicted binding affinity than the second, and we therefore hypothesized that its disruption (Ps1) would have a more pronounced effect on promoter activity than disruption of the second peak. To test this hypothesis, we synthesized (by GenScript) a 600 bp fragment of the IFIT2 promoter (chr10:89300997–89302495) and cloned it upstream of an eUnaG2 fluorescent reporter. This construct allowed us to assess promoter activity under various experimental conditions in life HeLa cells. WT and *IRF1* KO cells were transfected with wild-type and scrambled promoter-reporter constructs. Forty-eight hours post-transfection, fluorescence intensity was measured by flow cytometry. The experiment was conducted under three conditions: untreated, IRF1 OE, and IFN-β treatment for 12 hours. All conditions were tested for the four constructs: WT, Ps 1, Ps 2, and Ps 1+2. Transfecting WT HeLa cells with the WT promoter construct, showed significantly increased fluorescence either upon IRF1 OE or IFN-β treatment. This is consistent with increased transcription activity. In contrast, all three scrambled constructs exhibited reduced fluorescence intensity under these conditions, indicating disruption of IRF1-mediated promoter activation. Among the scrambled constructs, Ps 2 retained the highest activity, while Ps 1 and Ps 1+2 showed markedly reduced signal, aligning with our prediction that the first binding site is more important. In IRF1 KO cells, baseline fluorescence remained low across all constructs, and no induction was observed in the absence of stimulation. However, upon IFN-β treatment, the wild-type promoter again showed increased fluorescence, whereas the Ps 1 and Ps 1+2 variants remained less responsive. This further supports the conclusion that IRF1 binding is required for full promoter activation and that the predicted motifs are functionally important.

Together, these findings confirm that IRF1 directly regulates the IFIT2 promoter through specific sequence elements, and that our computational model reliably predicts functional IRF1 binding sites. By combining promoter mutagenesis, transcriptional reporter assays, and integrative modeling, we demonstrate that IRF1 binding is both necessary and sufficient for transcriptional activation of this key ISG under multiple signaling conditions. This approach not only validates our motif-based prediction strategy but also reinforces the central role of IRF1 as a versatile regulator of antiviral gene expression.

## Discussion

Our investigation into the role of IRF1 in regulating interferon responses has revealed its multifaceted and context-dependent functions in both basal and induced immune states. Traditionally viewed as an ISG, IRF1 is often characterized primarily by its role in amplifying responses to IFN stimulation. However, our findings challenge this restricted view, showing that IRF1 also plays a constitutive role in regulating immune homeostasis and antiviral readiness, even in the absence of cytokine stimulation. Contrary to our initial hypothesis (11) *IRF1* KO did not impair the induction of ISGs upon IFN-β stimulation (Fig. S1A), suggesting that the canonical JAK-STAT pathway can compensate for the absence of IRF1 under acute type I IFN signaling. This is consistent with reports showing that ISG induction is mediated by STAT1, STAT2, and IRF9-containing complexes, which bind ISRE and GAS motifs and contribute to redundant or overlapping regulatory networks (78–80). These complexes can bind both ISRE and GAS sites, leading to some redundancy in the signaling pathways. Therefore, the nuanced role of IRF1 in this process may reflect this functional overlap in immune responses (70). In line with this hypothesis, our RNA-seq data and WB analysis revealed increased mRNA and protein abundance of other IRFs, specifically IRF3, IRF7, and IRF9, in *IRF1* KO cells (Fig. S1C and E), suggesting potential compensatory transcriptional activity. Indeed, the IFIT2 promoter was activated both in the presence of IRF1 and upon its KO, to high levels (Fig. 8G).

Nevertheless, transcriptomic analysis revealed that IRF1 is critical for maintaining basal ISG expression under homeostatic conditions. *IRF1* KO cells exhibited substantial upregulation of antiviral genes such as *MX1*, *IFIT3*, and *IL6*, along with the downregulation of regulatory genes such as *AQP3* and *LPL* indicating a loss of transcriptional control that could prime cells for aberrant inflammatory responses (Fig. S1B) (81). Our pathway analysis using QIAGEN Ingenuity Pathway Analysis (IPA) revealed that IRF1 exerts a broad regulatory influence on immune signaling networks, functioning both as a transcriptional activator and repressor depending on gene context and signaling conditions. Without external stimulus, IRF1 suppresses key components of the innate immune system, including MAVS and type I/II interferon signaling pathways (Fig. 2D), while positively regulating other immune programs such as IL6/STAT3 signaling, which were downregulated in *IRF1* KO cells (Fig. 2E). These observations support a dual regulatory role for IRF1 in fine-tuning immune responses. Moreover, they are consistent with previous reports describing IRF1’s essential role in maintaining immune homeostasis, promoting basal gene expression, and preventing the aberrant activation of inflammatory and stress-related pathways (82, 83).

IRF1 OE triggered potent antiviral protection even in the absence of exogenous IFN treatment, indicating that IRF1 can activate an antiviral program autonomously. Indeed, RNA-seq showed IRF1 OE to induce gene transcription similar to that observed upon IFN-β treatment (but not IFN-γ treatment, Fig. S7A and B). Therefore, it is not surprising that this protection was strictly dependent on the presence of functional IFNARs in the recipient cells but was independent of both IFNARs and JAK1 in the IRF1-OE producer cells (Fig. 3D), suggesting that IRF1 can induce the secretion of antiviral factors that act in a paracrine manner. Notably, some antiviral protection persisted in *JAK1* KO IRF1 OE cells, as well as in cells treated with the pan-JAK inhibitor Ruxolitinib. This observation suggests that IRF1 can mediate antiviral responses through alternative signaling routes independent of classical JAK-STAT activation, potentially involving non-canonical pathways such as MAPK or PI3K/mTOR signaling. These findings are consistent with previous reports showing that IRF1 can directly upregulate antiviral effectors without requiring JAK1-mediated STAT phosphorylation, supporting the idea of a non-canonical IRF1-driven antiviral pathway (80, 84).

Transcriptomic analysis of IRF1 OE cells revealed robust upregulation of multiple type I interferon genes, particularly members of the IFN-α family, including IFNA1, IFNA2, and IFNA4 (Fig. 4C). This upregulation was also detected in *JAK1* KO and IFNAR KO backgrounds, indicating that while feedback suppression mechanisms may exist, they do not fully inhibit IRF1-mediated transcription. At the protein level, mass spectrometry of concentrated conditioned media confirmed the presence of secreted IFN-α peptides, identifying two unique peptides corresponding to IFN-α1 and one corresponding to IFN-α4 (Fig. S5A–B, S6A–B). These results support the hypothesis that IRF1 directly induces the secretion of type I IFNs, particularly IFN-α subtypes, which in turn are sufficient to establish paracrine antiviral protection in neighboring cells.

A challenge in our study was to accurately identifying specific IFN-α subtypes in complex biological samples. The high sequence similarity among IFN-α family members leads to overlapping peptide signatures, making it difficult to distinguish them confidently in mass spectrometry datasets. This is a well-documented limitation, as peptide fragments that align with multiple IFN-α isoforms are often excluded from analysis due to ambiguity. Similar challenges have been reported in the literature, highlighting the need for more refined detection tools such as immunoassays with minimal cross-reactivity to resolve and quantify individual IFN species (85). To address this issue, we employed the LEGENDplex™ Human Type 1/2/3 Interferon Panel (BioLegend, Cat. 741271), a multiplex bead-based immunoassay designed for flow cytometry. This assay enables simultaneous detection of multiple interferons including IFN-α2, IFN-β, IFN-γ, IFN-λ1, and IFN-λ2/3 with high specificity and sensitivity. Using this platform, we directly detected IFN-α2 protein in the conditioned media of IRF1-overexpressing cells (Fig. 4D–E, Fig. S4D–E), confirming IRF1-induced cytokine secretion at the protein level and validating our transcriptomic and mass spectrometry findings.

Another interesting observation emerged from the transcriptomic analysis of IRF1-overexpressing cells. After hierarchical clustering of the RNA-seq data, we compared IRF1 OE profiles to those of WT cells treated with IFN-β for 16 hours. This comparison was intended to isolate gene expression changes specific to IRF1 rather than those driven by general IFN signaling. Through this approach, we identified a distinct group of genes in cluster 3 that were upregulated following IRF1 overexpression but not in response to IFN-β treatment (Fig. 5A and Fig. S7C). Pathway enrichment analysis of cluster 3 using Ingenuity Pathway Analysis (IPA) revealed significant activation of pathways related to the adaptive immune system, particularly those involving T cell signaling and function (Fig. 5B). This finding aligns with previous studies linking IRF1 to adaptive immunity, including its role in T cell development and maturation (47, 48). These results suggest that IRF1 may extend its regulatory influence beyond antiviral and innate immune responses, contributing to the modulation of adaptive immune programs under specific conditions.

In addition to its role in activating immune-related genes, our findings also highlight IRF1’s ability to fine-tune gene expression by acting as both a transcriptional activator and repressor. RNA-seq analysis of *IRF1* KO and OE revealed distinct sets of upregulated and downregulated genes under each condition. Surprisingly, we also observed a subset of genes that were upregulated in both the knockout and overexpression conditions, as illustrated in the Venn diagram in Fig. 5C. This overlap suggests that IRF1 may exert opposing regulatory effects on certain genes depending on context potentially through secondary signaling effects or dosage-sensitive promoter interactions (86–89). Our data suggest that this may be due to the increased abundance of IRF3, IRF7 and IRF9 in IRF1 KO cells. Interestingly, IRF3 protein levels were elevated in IRF1 KO cells regardless of treatment, suggesting a constitutive upregulation. IRF7 abundance increased in WT cells after IFN-β stimulation, as anticipated. However, its abundance increased also in untreated *IRF1* KO cells. A similar trend was observed for IRF9. These results were corroborated by qPCR analysis, which mirrored the protein-level changes (Fig. S1C and E). IRF1, IRF3 and IRF7 can bind ISRE-like sequences. However, IRF1 also recognizes single IRF-E half-sites (GAAA), whereas IRF7 typically binds dimeric GAAA sites. Prior studies have shown that IRF7 overexpression is sufficient to drive ISG induction and confer antiviral protection (90). Moreover, the highly increased abundance of IRF9 can also stimulate some ISG gene expression without type I IFN stimulation(91). Thus, the partial redundancy between the different IRFs can explain why some ISGs are activated in IRF1 KO cells. Our data support a model in which IRF1 suppresses other IRFs under homeostatic conditions directly or indirectly. In its absence, IRF7, IRF3 and IRF9) become derepressed, contributing to the elevated ISG expression and partial antiviral resistance observed in IRF1 KO cells.

PBM experiments were used to identify specific DNA binding motifs for IRF1, which were used to establish a predictive binding affinity model (Fig. 7, S10, and S11). The integration of our predictive binding affinity model with the reprocessed raw ChIP-seq data provides significant insights into the role of IRF1 in regulating gene expression through precise promoter interactions. By calculating binding affinities for 7-mer sequences and focusing on promoter regions of genes with known regulation patterns from RNA-seq analysis, our model successfully identified high-affinity binding sites for IRF1. The alignment of these predicted sites with the ChIP-seq data underscores the accuracy and robustness of our approach in mapping key regulatory interactions. A particularly notable finding was the identification of a small but discernible peak in the IFN promoter region. Although this peak was initially underrepresented in the ChIP-seq data, our affinity-based model highlighted it as a potential binding site. This subtle signal is consistent with the biological context, as IFN receptors exhibit high sensitivity for their ligands. This observation underscores the capacity of our model to detect binding events that, while subtle, play a crucial role in initiating immune responses. Moreover, the correlation between regions of strong ChIP-seq coverage and our predicted high-affinity sites further supports the utility of our model in pinpointing critical binding locations for IRF1. We validated our binding predictions using the IFIT2 promoter as a case study. This gene was upregulated in both *IRF1* KO and OE cells and served as a clean model to dissect promoter dynamics. By scrambling predicted IRF1 binding motifs and measuring eUnaG2 reporter expression, we showed that mutation of either binding site impaired promoter activity with the first site having the strongest predicted and functional impact (Fig. 8). These results demonstrate that IRF1 binding is both necessary and sufficient for driving transcription from target promoters, and that our computational framework can successfully identify biologically relevant regulatory elements. To explore potential compensatory mechanisms underlying residual antiviral activity in IRF1 KO cells, we extended our promoter analysis to IRF3, IRF7, and IRF9. key members of the IRF family that are upregulated upon IRF1 KO (Fig. S1C). Using our PBM-derived predictive binding model and reprocessed raw ChIP-seq data we examined the promoter regions of these genes and found strong predicted IRF1 binding affinity as well as ChIP-seq enrichment (Fig. S12A–C), suggesting that IRF1 may directly regulate their transcription.

In conclusion, IRF1 emerges as a multifaceted regulator of immune responses, playing essential roles in both immune homeostasis and antiviral defense. Our study demonstrates that IRF1 not only orchestrates the secretion of key cytokines particularly type I IFNs but also fine-tunes the expression of immune-related genes through a dual function as both a transcriptional activator and repressor. This regulatory versatility allows IRF1 to maintain basal expression of ISGs and coordinate antiviral defenses, even in the absence of canonical JAK-STAT signaling. Importantly, the partial antiviral protection observed in JAK1 knockout cells highlights IRF1’s capacity to engage non-canonical signaling pathways potentially involving MAPK or PI3K/mTOR cascades thereby extending its functional reach beyond traditional IFN responses. These findings are particularly relevant in the context of cancer immunotherapy, where defects in JAK1 signaling can limit the efficacy of cytokine-based treatments (92–94). By driving innate immune programs independently of JAK-STAT activation, IRF1 represents a promising therapeutic target for enhancing antiviral and antitumor immunity in settings of signaling impairment. The successful application of our affinity-based predictive model further underscores the value of integrating computational and experimental approaches. This strategy enabled the identification of both dominant and subtle IRF1 binding events that may be overlooked by traditional ChIP-seq analysis, providing a more nuanced understanding of IRF1’s regulatory landscape. Our findings position IRF1 as a central node in interferon biology balancing gene activation and repression, modulating cytokine output, and initiating robust antiviral programs independently of upstream signaling fidelity.

Future studies should investigate how IRF1 cooperates with other transcription factors such as IRF3, IRF7, IRF9, STAT1, and STAT2 to coordinate gene expression networks, and whether IRF1 binding affinity or chromatin context governs its context-specific activity. The ability of IRF1 to selectively induce IFN-α subtypes also opens exciting therapeutic avenues, particularly in immuno-oncology, where restoring or mimicking IRF1 function may reinstate immune competence in tumors with impaired upstream pathways.

### Experimental Procedures

#### Cell lines

HeLa cells, derived from a human cervical cancer cell line, were used for all experiments in this study and were cultured in Dulbecco’s Modified Eagle’s Medium (DMEM, Gibco 41965-039) supplemented with 10% fetal bovine serum (FBS, Gibco 12657-029), 1% pyruvate (Biological Industries 03-042-1B), and 1% penicillin-streptomycin (Biological Industries 03-031-1B).

#### Generation of KO cells with CRISPR-Cas9

All knockout (KO) cell lines in this study were generated with CRISPR-Cas9 technology, as previously described (11, 95). For the current study, we generated *IRF1* KO HeLa cell lines. To target the appropriate genes, we designed a single guide RNA (sgRNA) with the Benchling CRISPR Design Tool. The sgRNA sequence (5’-GATGCTTCCACCTCTCACCA-3’) was designed to target exon 3 of the *IRF1* gene. The sgRNA was subcloned into the pX459 plasmid (Addgene plasmid #62988) and used to transfect HeLa cells with JetPRIME (Polyplus 114-07) according to the manufacturer’s instructions. Clones were selected using puromycin resistance, and single cells were expended and verified by Western blotting analysis and genomic sequencing for the KO. Similarly, *JAK1* KO and double *IFNAR1/IFNAR2* KO cell lines were generated as previously described (11, 12, 95).

#### IRF1 and GFP Over expression

Transient transfections of HeLa cells were performed with JetPRIME reagent (Polyplus 114-07) according to the manufacturer’s protocol. Forty-eight hours later, the cells were treated to assess antiviral activity and gene expression, and protein phosphorylation was determined by Western blotting analysis. Constructs Used: IRF1: Full-length protein inserts in mammalian vector pDisplay. GFP: Full-length protein inserts in mammalian vector pDisplay.

#### Western blotting analysis

For the Western blot analysis, cells were lysed in PBS (pH 7.4) supplemented with 1% NP-40, 1 mM EDTA, and a mix of protease inhibitors (Sigma P8340), phosphatase inhibitor cocktail 2 (Sigma P5726), and phosphatase inhibitor cocktail 3 (Sigma P0044). The lysates were separated using 4 to 20% SDS-PAGE (GenScript M00657) and transferred onto a 0.45 µm nitrocellulose membrane (Bio-Rad). Membranes were blocked with 5% bovine serum albumin (BSA) before primary antibody incubation. Detection of specific protein bands was carried out using enhanced chemiluminescence (ECL) substrate. The primary and secondary antibodies used are listed in Table S2, including HRP-conjugated anti-mouse (Jackson ImmunoResearch, 115035146) and HRP-conjugated anti-rabbit (Jackson ImmunoResearch, 111035144). The Western blots were visualized using the Odyssey Fc imaging system (LI-COR Biosciences). Band intensities were quantified with Image Studio Lite software (LI-COR Biosciences). For normalization, the following steps were applied: (1) The highest signal intensity band on each membrane was set as the reference with a value of 1. (2) Other bands on the same membrane were normalized by dividing their intensities by this reference value. (3) To normalize across multiple membranes, each band’s intensity detected by a particular antibody in each experiment was further divided by the normalized value of the loading control, tubulin. (4) For phosphorylated proteins, the intensity of each band was additionally normalized by dividing the phosphorylated protein’s intensity by the intensity of the corresponding total protein.

#### Antiviral and antiproliferative assays

HeLa cells (1.2 × 10^4^ cells for antiviral assays and 2 × 10^3^ cells for antiproliferative assays) were seeded into flat-bottomed 96-well plates and cultured overnight. For both assays, cells were treated with ten three-fold serial dilutions of either IFN-β or IFN-γ. For IFN-β, the starting concentrations were 500 pM for the antiviral assay and 50 nM for the antiproliferative assay, with cells incubated for 4 hours prior to virus addition. For IFN-γ, the starting concentrations were 500 pM for the antiviral assay and 50 nM for the antiproliferative assay, with an 8-hour incubation before virus addition in the antiviral assay. Antiviral protection against VSV (vesicular stomatitis virus) and EMCV (encephalomyocarditis virus) was evaluated by assessing the inhibition of virus-induced cytopathic effects. After the respective incubation periods with interferons, VSV or EMCV were introduced into the wells, followed by an 18-hour incubation for VSV or 20 hours for EMCV. Antiproliferative activity was measured 96 hours post-treatment with either IFN-β or IFN-γ. Cell viability was determined through crystal violet staining in a 96-well plate format, with absorbance readings at 590 nm using a TECAN INFINITE M PLEX plate reader, capturing 5×5 multiple reads per well. Data normalization was performed using the formula:

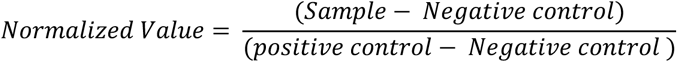

where “Sample” refers to treated cells exposed to the virus, “Negative Control” refers to cells exposed to the virus without treatment, and “Positive Control” refers to untreated cells without virus exposure. EC50 values and cell sensitivity to the treatments were calculated by fitting the response curves using GraphPad Prism software (version 9.5.0.730).

#### Intrinsic antiviral protection

HeLa cells (2 × 10⁴) were seeded into flat-bottomed 96-well plates and cultured overnight. To assess the cells’ intrinsic antiviral capacity, either VSV or EMCV was added directly to the wells without any prior treatment. Cells were incubated with VSV for 8 hours or EMCV for 12 hours. Following incubation, cell viability was determined by crystal violet staining, as described in the antiviral and antiproliferative assay section. Absorbance was measured at 590 nm using a TECAN INFINITE M PLEX plate reader, and data were normalized accordingly.

#### Quantitative PCR analysis

The relative expression levels of selected human interferon-stimulated genes (ISGs) were determined using the Applied Biosystems ViiA 7 Real-Time PCR System, following previously established protocols (5, 11, 95). PCR reactions were performed using Fast SYBR Green Master Mix (Applied Biosystems), and cDNA was synthesized from 1 μg of total RNA using the high-capacity cDNA reverse transcription kit (Applied Biosystems). Total RNA was extracted with the NucleoSpin RNA kit (Macherey-Nagel). For qPCR, 14.5 ng of cDNA was used per reaction, with a total reaction volume of 7 μl, using the appropriate primers listed in Table S1. Relative expression levels were calculated using the ΔΔCT (cycle threshold) method, with the fold-change expression determined using the formula RQ = 2^−ΔΔCT^. Hypoxanthine-guanine phosphoribosyltransferase 1 (*HPRT1*) served as the reference gene for normalization across samples.

#### RNA-seq Sample Preparation, Library Preparation, and Sequencing

RNA-seq analysis was performed on HeLa wild-type (WT) cells, *IRF1* KO cells, and cells with overexpression of IRF1 or GFP as a control. All the samples were done in two biological duplicates. The specific conditions included:

WT HeLa cells (untreated) for basal RNA expression.

WT HeLa cells treated with 2 nM IFN-β for 16 hours.

WT HeLa cells treated with 100 nM IFN-γ for 6 and 16 hours.

*IRF1* KO cells (untreated).

*IRF1* KO cells treated with 100 nM IFN-γ for 6 and 16 hours.

WT HeLa cells overexpressing IRF1.

*JAK1* KO cells overexpressing IRF1.

*IFNAR*s KO cells overexpressing IRF1.

WT HeLa cells overexpressing GFP as a control for transfection.

RNA was isolated as described for qPCR analysis. Library preparation and RNA-seq experiments were conducted at the INCPM units at the Weizmann Institute of Science. Samples were sequenced using the Illumina NovaSeq SP platform with 100-cycle runs. Library preparation included the addition of Unique Molecular Identifiers (UMIs) to each DNA fragment during reverse transcription using an oligo dT primer. UMIs are short molecular tags that help to reduce errors and quantitative bias during PCR amplification by uniquely labeling each original RNA molecule before amplification, following protocols described in previous studies (96, 97).

#### DNA-seq data analyze

RNA-seq data were analyzed using the UTAP (Universal Transcriptome Analysis Pipeline), a robust pipeline developed by the Weizmann Institute of Science for high-throughput transcriptomic analysis (55), The UTAP pipeline performs a comprehensive workflow, including quality control, read alignment, transcript quantification, and differential expression analysis. Raw sequencing reads were first assessed for quality using FastQC to identify any issues in read quality, adapter content, or sequence duplication levels. Reads were then aligned to the human genome reference (hg38) using STAR (98), a highly efficient RNA-seq aligner. The UTAP pipeline automatically handles the alignment process, mapping reads to annotated gene models and providing high-quality alignments for subsequent analysis. Differentially expressed genes (DEGs) were identified and normalized using DESeq2 (56). Three thresholds were applied to determine significant DEGs: (1) an adjusted p-value (p-adj) of ≤ 0.05, (2) a log2 fold change of ≥ 2 or ≤ -2, and (3) a base mean value above 5 to ensure that all samples exhibited a minimum level of expression.

#### Heatmap Generation and Clustering Analysis

RNA-seq data were analyzed to identify expression patterns and classify genes into distinct clusters. Hierarchical clustering and heatmap visualization were performed using Python libraries, including pandas (99), numpy (100), scipy (101), scikit-learn (102), matplotlib (103), and seaborn (104). RNA-seq data were subjected to hierarchical clustering using Ward’s method, implemented with the linkage function from the scipy library. A dendrogram was generated to visualize the clustering relationships among samples, indicating the similarity of expression profiles. Based on the dendrogram, a flat clustering was performed using the fcluster function, dividing the data into four clusters. Heatmap of Median Expression with Annotations: For each identified cluster, the median expression, interquartile range (IQR), and standard deviation were calculated. These values were visualized in a heatmap with cluster-specific annotations, displaying median expression values alongside their respective IQR and standard deviation. Cluster labels included the number of genes in each cluster, providing additional context.

#### Pathway Analysis Using QIAGEN Ingenuity Pathway Analysis (IPA)

To interpret the biological significance of RNA-seq results, we performed pathway analysis using QIAGEN Ingenuity Pathway Analysis (IPA) (57). Genes from the RNA-seq clustering analysis were uploaded to the IPA software, where they were mapped to known molecular networks and pathways. The analysis focused on identifying pathways, upstream regulators, and interaction networks associated with differentially expressed genes. For inclusion in the IPA analysis, genes were selected based on a threshold of an adjusted p-value ≤ 0.05 and a log_2_ fold change ≥ 2 or ≤ -2, as described in the IPA user guide. These criteria ensured that only genes with statistically significant changes in expression and meaningful fold changes were considered in the analysis. The IPA software generated a Graphical Summary, which provides an overview of key pathways, predicted regulators, and molecular interactions derived from the RNA-seq data.

#### LEGENDplex 5-Plex Cytokine Assay

In this study, we used the LEGENDplex™ 5-Plex kit (Cat. 741271) to measure cytokine levels in conditioned media, following the manufacturer’s protocol (64). Conditioned media were collected from target cells 48 hours post-transfection and incubated with cytokine-specific capture beads for 1 hour. After incubation, samples were washed and analyzed using flow cytometry (Cytoflex S, Beckman Coulter). The data obtained from the flow cytometry analysis were processed using the LEGENDplex™ software, which facilitated quantification of the cytokines. Appropriate controls, as specified by the manufacturer, were included in each assay to ensure accurate and reliable results.

#### Preparation and analysis of conditioned medium

Conditioned medium (CM) from WT and *IFNAR*s KO cells. 17 ml of CM was collected from each cell type. The CM was concentrated using an Amicon Ultra centrifugal filter unit with a 10 kDa molecular weight cutoff, reducing the volume to 500 μl. The concentrated CM was then subjected to size-exclusion chromatography using a Superdex® 200 10/300 GL gel filtration column (GE, cat. 28-990944). Fractions of 1 ml each were collected, and the antiviral activity of each fraction was tested for its ability to protect against vesicular stomatitis virus (VSV) infection, using a protocol identical to that employed for IFN-β, except for the use of CM as the treatment.

#### Mass Spectrometry

Samples were digested with trypsin overnight at 37°C and analyzed using a Vanquish liquid chromatography system (Thermo Scientific) coupled to a timsTOF SCP mass spectrometer equipped with a CaptiveSpray ion source (Bruker Daltonics). The mass spectrometer was operated in positive data-dependent acquisition mode. One microliter of the peptide mixture was injected by an autosampler onto a C18 trap column (Pepmap Neo C18, 5 µm, 0.3 x 5 mm, Thermo Scientific). After trapping, peptides were eluted from the trap column and separated on a C18 analytical column (Pepsep C18, 150 x 0.15 mm, 1.5 µm, Bruker Daltonics) using a linear gradient of 5% (v/v) to 35% (v/v) acetonitrile in water over 35 minutes, with a flow rate of 1.5 µL/min. Both the trap and analytical columns were maintained at 50°C. The timsTOF SCP was configured using standard proteomics PASEF (Parallel Accumulation-Serial Fragmentation) parameters. The target intensity per individual PASEF precursor was set to 20,000, with an intensity threshold of 1,500. The ion mobility scan range was set between 0.6 and 1.6 V·s/cm² with a ramp time of 100 ms, and 10 PASEF MS/MS scans were acquired per cycle. Precursor ions in the m/z range of 100 to 1700, with charge states between 2+ and 6+, were selected for fragmentation. Active exclusion was enabled for 0.4 minutes. Raw data were processed using PeaksStudio 10.0 software (Bioinformatics Solutions, Canada). Search parameters included trypsin (semi-specific) as the enzyme, with carbamidomethylation set as a fixed modification and oxidation of methionine and acetylation of protein N-terminus as variable modifications. The data were searched against a database containing interferon sequences.

#### IRF1 DBD-mNeonGreen

The expression of IRF1 DBD-mNeonGreen was achieved by introducing the plasmid pET28-*IRF1* DBD-mNeonGreen into T7 competent bacterial cells. The transformed bacteria were cultured in 2YT media supplemented with kanamycin for selection. After growth, the bacterial cells were harvested and lysed using sonication in PBS buffer containing 360 mM NaCl, 10% glycerol, protease inhibitors, and DTT to prevent protein degradation. Following lysis, the lysate was clarified by centrifugation and further filtered using a 0.45 µm filter. The filtered lysate was then loaded onto a Hi-Trap SP HP anion exchange column (GE Healthcare, Cat. 17115101) for initial purification. The final purification step involved size exclusion chromatography using a Superdex 75 Increase 16/600 column (GE Healthcare, Cat. 28-989333) to ensure complete purification and separation of the IRF1 DBD-mNeonGreen protein.

After protein purification, three different samples were prepared to assess DNA binding specificity. The first sample contained purified protein without any DNA. The second sample included the purified protein incubated with a synthesized DNA oligonucleotide containing a specific IRF1 binding sequence (5’-GAGAAGTGAAAGTACTTTCACTTCTC-3’) at a 1:1 molar ratio of protein to DNA. The third sample consisted of the purified protein with a scrambled control DNA sequence (5’-ATATTACGTCGCACTAGGATAGATCT-3’), also added at a 1:1 molar ratio. All samples were incubated for 4 hours at room temperature before measurement to ensure sufficient binding interaction. These samples were then used to evaluate the protein’s specific binding affinity and interaction with DNA in subsequent assays.

#### Size-Exclusion Chromatography (SEC)

A sample was manually injected onto a Superdex 200 Increase 10/300 GL column (GE, cat. 28-990944). The column was pre-equilibrated with PBS at pH 7.4, supplemented with 360 mM NaCl and DTT. To determine the molecular mass of the eluted proteins, a standard curve from a previously published study (105) was used.

#### Mass Photometry

Microscope coverslips (no. 1.5, 24 × 50, cat# 0107222, Marienfeld) were cleaned by sequential sonication in 50% isopropanol (HPLC grade)/Milli-Q water and Milli-Q water alone for 5 minutes each, followed by drying with a nitrogen stream. Four gaskets (Reusable culturewell™ gaskets, 3 mm diameter × 1 mm depth, cat# GBL103250-10 EA, Sigma-Aldrich) were cut into a 2 × 2 array, cleaned using the same method as the coverslips, and placed on top of the coverslip, with each well used for a separate sample measurement. Immediately before mass photometry measurements, protein stocks were diluted in PBS at pH 7.4. A fresh buffer was added to each well to find the focal position, which was then secured using an autofocus system based on total internal reflection for consistent focus throughout the measurements. For each measurement, 5 μl of diluted protein solution (at nanomolar concentrations) was added to the well, and after the autofocus system stabilized, 120-second movies were recorded. Each sample was measured in triplicate. Calibration of the contrast-to-mass conversion was performed in the same measurement buffer using urease (Sigma cat# U7752-1VL) as a reference, with known oligomer masses. All data were acquired using a OneMP mass photometer (Refeyn Ltd, Oxford, UK). Data collection was done using AcquireMP software (v2.2, Refeyn Ltd), and data analysis was performed with DiscoverMP software (v2.3.0, Refeyn Ltd).

#### Nano DSF melting assay

Protein thermal stability was assessed using the Tycho NT.6 instrument (NanoTemper) (106). Protein samples were prepared at a concentration of 0.1 to 0.5 mg/ml in PBS. A total of 10 µl of each protein sample was loaded into Tycho NT.6 capillaries and inserted into the instrument. Thermal unfolding was monitored by heating the samples from 35°C to 95°C while recording fluorescence at 330 nm and 350 nm. The fluorescence ratio (350/330 nm) was used to generate unfolding curves, and the melting temperature (Tm) was determined as the inflection point of the curve, indicating the temperature at which protein unfolding occurred.

#### Protein Binding Microarrays (PBM)

Experiments were conducted as previously described (67, 68). Briefly, custom-designed DNA microarrays (Agilent Technologies) were used, comprising ∼15,000 unique single-stranded 60-base oligonucleotides. The DNA library used here consists of de-Bruijn sequences representing all possible 9-mers and a constant 3’ end. The arrays were double stranded through incubation with a primer (complementary to the 3’ end constant region), Thermo Sequenase™ DNA polymerase, dNTPs, and 10X reaction buffer (Cytiva) for 2 hours at a temperature gradient (85°C to 60°C). The double-stranded microarray was pre-wet in PBS 0.01% Triton X-100 for 5 minutes and blocked with nonfat milk 2% (w/v) for 1 hour. Then, it was washed with PBS 0.1% Tween-20 for 5 minutes with PBS 0.01% Triton X-100 for 2 minutes. Purified IRF1-mNeonGreen was diluted to a final concentration of 2 μM in a binding mixture (nonfat milk 2%, 51.3 ng/μl salmon testes DNA, 0.2 mg/ml bovine serum albumin, 0.2% Triton X-100, PBS, 360 mM NaCl and 1 mM DTT). The microarray was incubated for 1 hour with the binding mixture at room temperature and then washed with PBS 0.5% Tween-20 for 3 minutes and with PBS 0.01% Triton X-100 for 2 minutes. IRF1-mNeonGreen fluorescence was scanned at 488 nm using a GenePix 4400A microarray scanner and the GenePix pro analysis software. For data analysis, signal was first spatially detrended, and position weighted matrices (PWMs) were generated using the “seed and wobble” algorithm (67). The energy-normalized logo for IRF1-mNeonGreen was generated using enoLOGOS (69).

#### ChIP-seq Data Analysis

ChIP-seq raw data (GEO accession number: GSE6928615 and GSE6928616) were reprocessed with a standardized pipeline using the Galaxy (71) platform following the methodologies outlined in (70). Data processing included quality control, read alignment, peak calling, and visualization. Raw reads were aligned to the human genome (hg38) using Bowtie2, followed by peak calling with MACS2 to identify regions of enriched IRF1 binding. Visualization of ChIP-seq coverage and comparison with predicted binding sites was performed using the Integrated Genome Browser (IGB) (72). Identified peaks were cross-referenced with predicted binding affinity scores to confirm potential IRF1 binding sites across the promoter regions of genes identified from RNA-seq analysis.

#### Generation of Predicted Binding Scores and BedGraph Files

To predict IRF1 binding affinity across promoter regions, we used a custom Python script to calculate z-scores for each 7-mer sequence, based on data from a UniProbe (107) binding affinity file generated from our PBM experiments. The DNA sequence of interest was defined, and the UniProbeZScoreFile parser was used to calculate binding affinity scores for each 7-mer, resulting in a vector of predicted binding scores. Genomic coordinates were assigned according to the target region. A BedGraph file was generated using a custom function, which iteratively mapped each score to its corresponding genomic position with a defined step size, creating a continuous profile of predicted binding affinity across the region. The resulting BedGraph file was used for visualization and comparison with ChIP-seq data in downstream analyses.

#### Visualization in the Integrated Genome Browser (IGB)

The generated BedGraph files containing predicted IRF1 binding scores were visualized using Integrated Genome Browser (IGB) (72). ChIP-seq data (GEO accession numbers: GSM6928615, GSM6928616) were loaded alongside the BedGraph tracks to allow direct comparison of predicted binding sites with observed ChIP-seq coverage. The genomic coordinates of each track were adjusted to ensure alignment, facilitating the identification of regions where predicted high-affinity binding sites overlapped with strong ChIP-seq peaks. To aid interpretability, predicted binding scores were transformed using an inverse log₂ function (2^z^-score) within IGB, restoring the relative binding signal to linear scale for more intuitive visualization. This visualization approach enabled a detailed comparison of computational predictions with experimental data, highlighting areas of potential IRF1 binding in the promoter regions of target genes.

## Data Availability

The RNA sequencing data generated in this study have been deposited in the NCBI GEO database and are accessible under the SuperSeries accession number PRJNA1328237.

## Acknowledgments

We gratefully acknowledge Moshe Goldsmith for invaluable assistance and for dedicated maintenance of the mass photometry instrument at the Department of Biomolecular Sciences, Weizmann Institute of Science.

## Authors Contributions

E. Z. investigation and data analysis; I. M., A. A. PBM measurements and analysis; S. B-D., D. H. RNA-seq analysis; J. Z. Mass spectrometry; E. Z., G. S. writing & editing; G. S., J. Z. funding acquisition; E. Z., G. S., conceptualization; G. S. supervision;

## Funding and additional information

This work was supported by a grant from Minerva Foundation number 714144, Israel Cancer Research Foundation number 1344142 and Israel Science Foundation number 3031/25. We acknowledge Structural mass spectrometry core facility of CIISB, Instruct-CZ Centre, supported by MEYS CR (LM2023042) and European Regional Development Fund-Project „UP CIISB“ (No. CZ.02.1.01/0.0/0.0/18_046/0015974) and the project National Institute of virology and bacteriology (Programme EXCELES, ID Project No. LX22NPO5103) - Funded by the European Union - Next Generation EU.

## Supporting Information

**Table S1.**
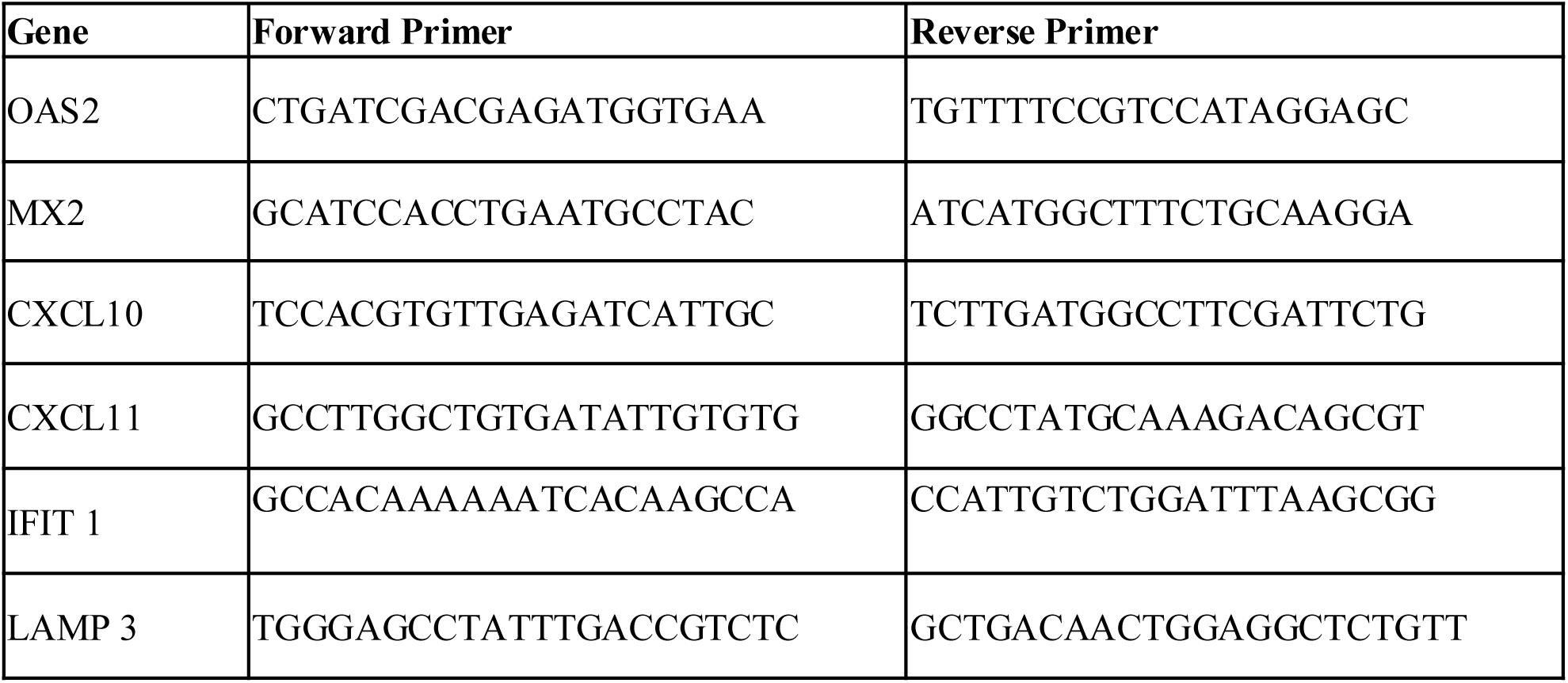
Primers used for real-time PCR.

**Table S2.** Primary antibodies for western blots.

| **Protein** | **Epitope** | **Concentration** | **Source** | **Catalogue No.** |
| --- | --- | --- | --- | --- |
| STAT1 | Total | 1:1000 | Cell Signalling Technology (CST) | 14995S |
| pSTAT1 | Tyr701 | 1:1000 | CST | 9167S |
| STAT2 | Total | 1:1000 | Santa Cruz Biotechnology | sc-1668 |
| pSTAT2 | pTyr690 | 1:1000 | CST | 4441S |
| IRF1 | Total | 1:1000 | CST | 8478S |
| Tubulin | Total | 1:5000 | Sigma-Aldrich | T9026 |

**Figure S1.**
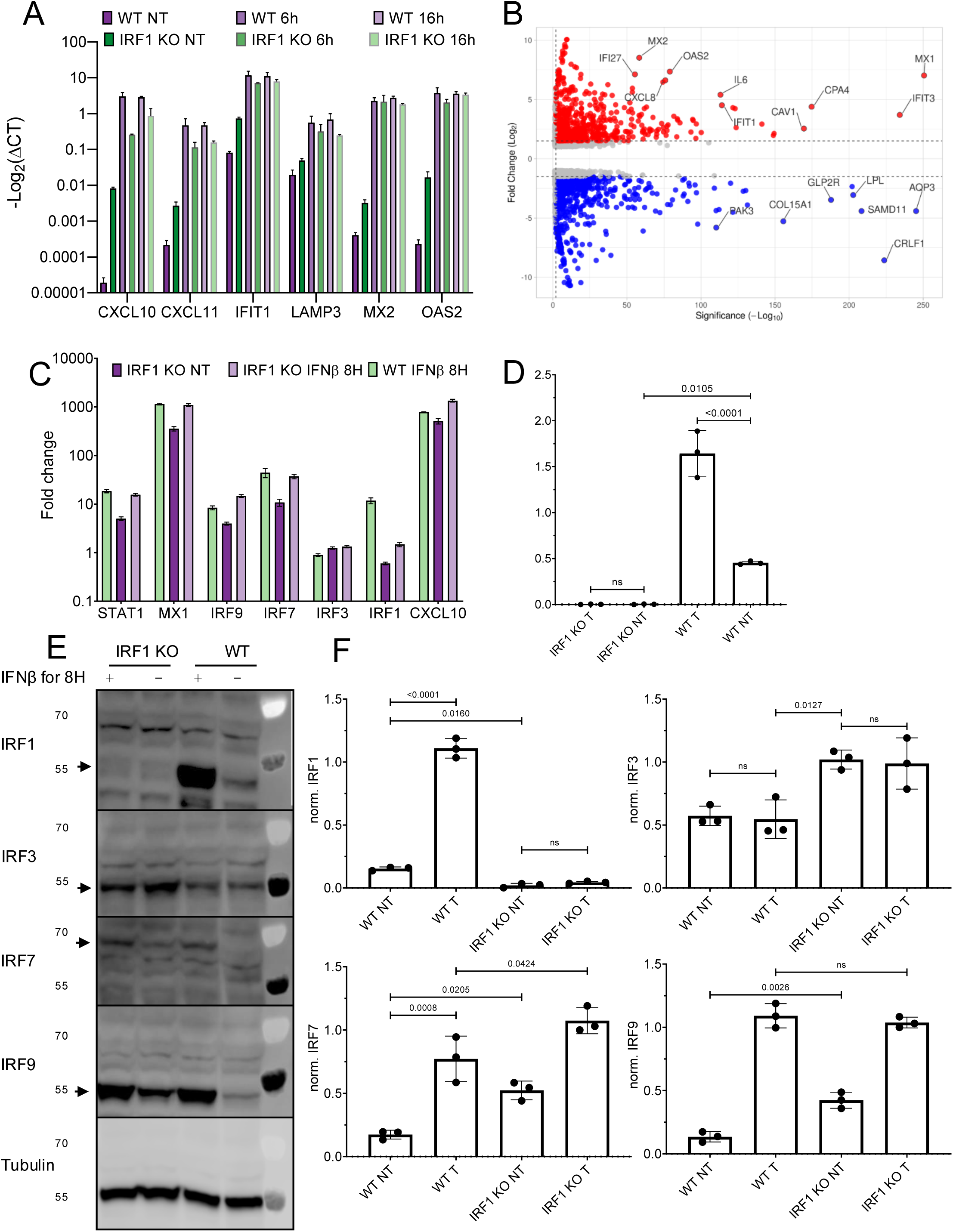
Basal gene expression abundance in *IRF1* KO cells. **(A)** RNA was extracted from WT and *IRF1* KO cells treated with IFN-β for 6 and 16 hours. The relative abundance of the indicated genes were determined by qPCR and are presented as -log2(ΔCT) normalized to that of *HPRT1. XAF1, MX1, MX2*, and *OAS2* are type I IFN induced robust genes; *IDO1, CXCL10* and *CXCL11* are type I IFN induced tunable genes. Data are means + SD of three independent experiments per group. **(B)** Volcano plot from RNA-seq result of *IRF1* KO compared to HeLa WT cells. The Y-axis is log_2_ fold change of *IRF1* KO non-treated over WT non-treated and the X-axis is -log_10_ of the Padjusted values. The threshold is 2>FC<-2. (**C**) WT and *IRF1* KO HeLa cells were left untreated or treated with IFN-β for 8 hours, and RNA was extracted for gene expression analysis. The relative mRNA levels of *STAT1, MX1, IRF1, IRF3, IRF7, IRF9*, and *CXCL10* were determined by qPCR. Data are presented as fold change compared with the expression in untreated WT cells and normalized to *HPRT1*. Data are means ± SD of three independent experiments per group. **(D**) Normalized total IRF1 abundance from the WB shown in Fig. 1A (and two more replicates). (**E**) WT HeLa cells and *IRF1* KO cells were left untreated or treated with IFN-β for 8 hours, and whole-cell lysates were analyzed by WB to detect IRF1, IRF3, IRF7, and IRF9 protein abundance. Tubulin was used as a loading control. Data are representative of three independent experiments. (**F**) Quantification of the Western blot signals shown in (E). Band intensities were normalized to tubulin and expressed relative to untreated WT cells.

**Figure S2.**
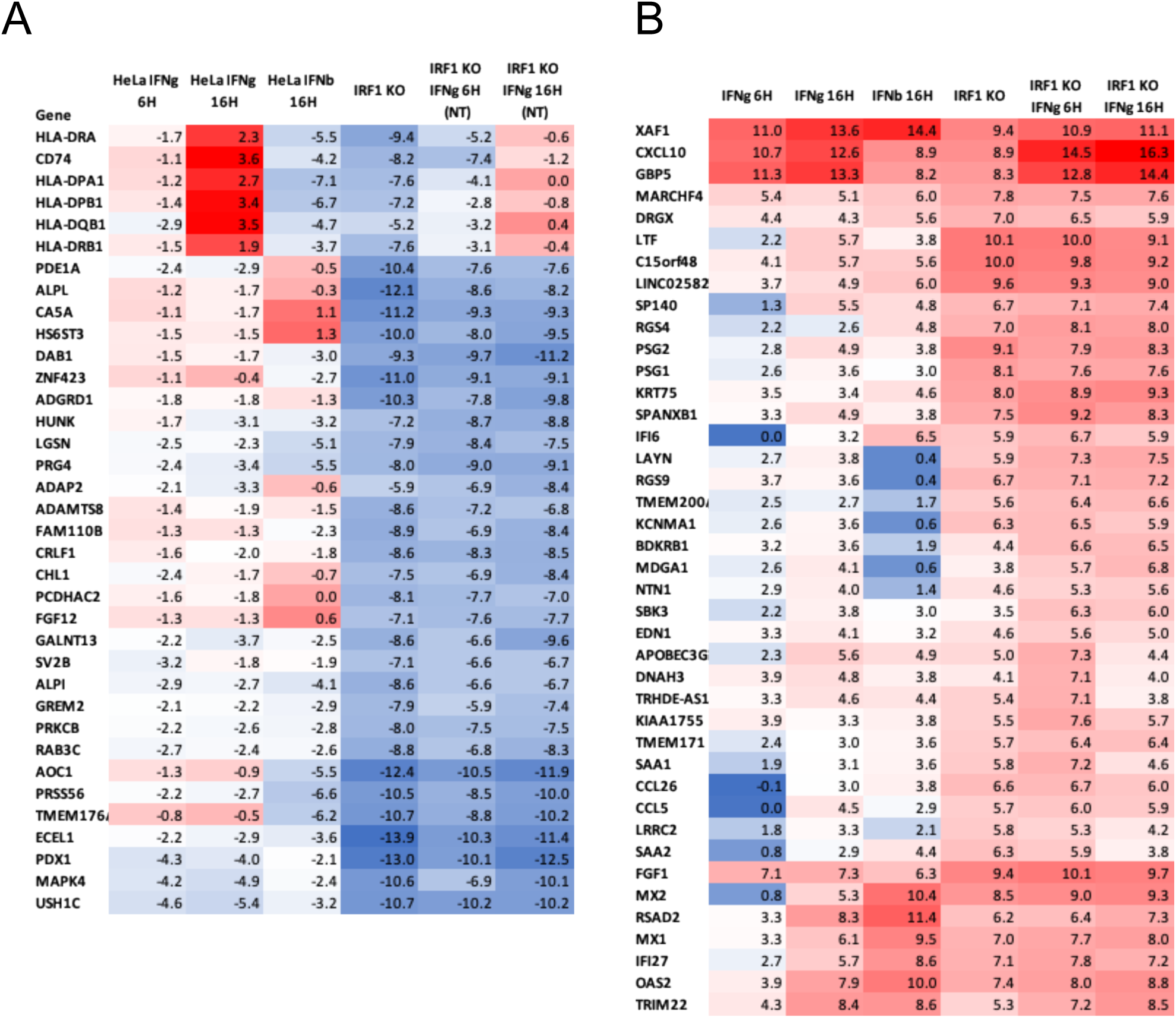
Gene expression changes in WT and *IRF1* KO cells. (A-B) Heatmaps displaying hierarchical clustering of genes from cluster 1 **(A)** and cluster 4 **(B)** from Fig. 2C, under the following conditions: WT cells treated with IFN-γ (100 nM) for 6 and 16 hours, WT cells treated with IFN-β (2 nM) for 16 hours, *IRF1* KO cells, *IRF1* KO cells treated with IFN-γ (100 nM) for 6 and 16 hrs. The values are –log(2) relative (to WT not treated) normalized counts from the RNA-seq data. In **(A)** the color scale is blue – lowest, white -2, red – highest. In **(B)** blue – lowest, white 3, red – highest.

**Figure S3.**
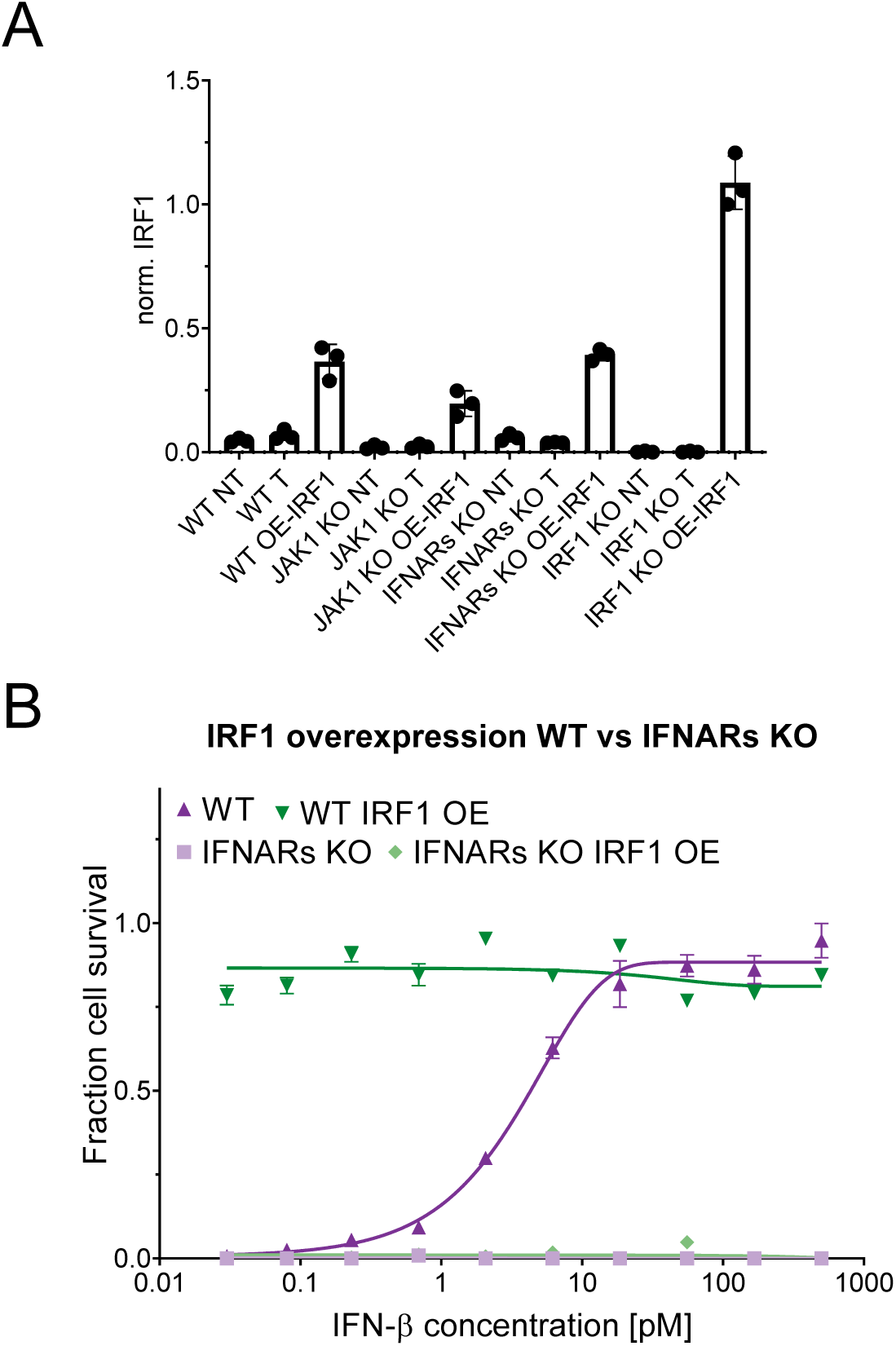
Impact of IRF1 overexpression on STAT Signaling and antiviral response in different KO cell lines. **(A**) Normalized total IRF1 abundance from WB shown in Fig. 3A and two more replicates**. (B)** Antiviral activity against VSV for WT HeLa, *IFNAR* KO and IRF1 OE cells 48 hrs post transient transfection of IRF1. Cells were treated with IFN-β for 4 hrs before infection with the VSV for 18 hrs. Cells were stained with crystal violet for cell viability. Data points are median of 3 independent experiments. Error bars represent the SD.

**Figure S4.**
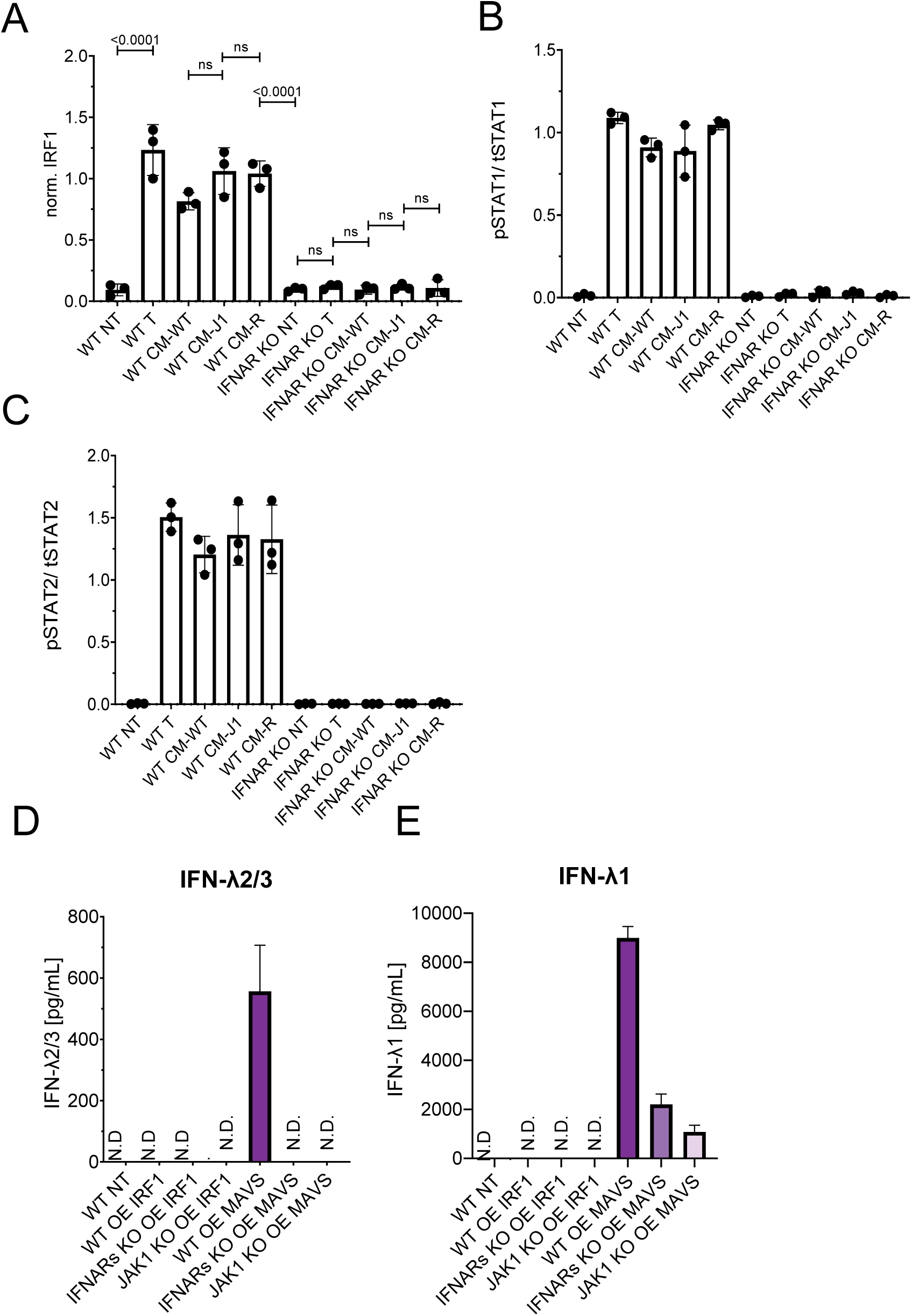
Quantification of cytokine signaling and flow cytometry analysis of IFN abundance in IRF1 and MAVS OE cells. **(A-C)** Quantification of WB shown in Fig. 4. **(A)** normalized pSTAT1 abundance relative to total STAT. **(B)** Normalized pSTAT2 abundance relative to total STAT2. **(C)** Normalized IRF1 abundance**. (D-E)** Flow cytometry analysis of IFN-λ2/3 **(D)** and IFN-λ1 **(E)** abundance in cells overexpressing IRF1 (IRF1 OE) or MAVS (MAVS OE) in WT, *JAK1* KO, or *IFNAR* KO cells. WT untreated (WT NT) cells were used as a negative control. The data represent the mean +SD from three independent experiments.

**Figure S5.**
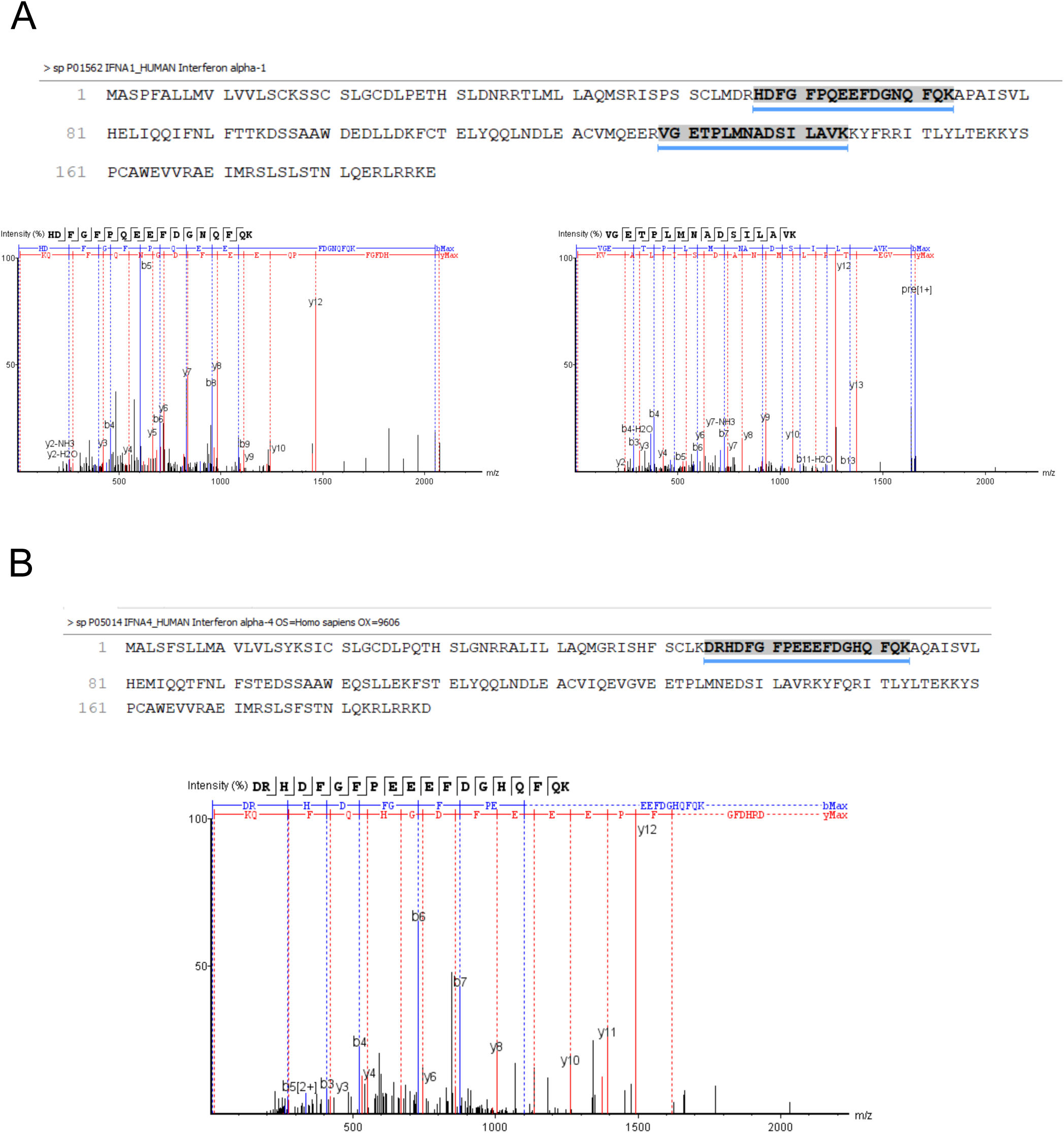
Mass spectrometry profile of protein fractions purified from conditioned media of WT cells. **(A)** Two unique peptides were identified as belonging to IFNα1. **(B)** One unique peptide was identified as belonging to IFNα4.

**Figure S6.**
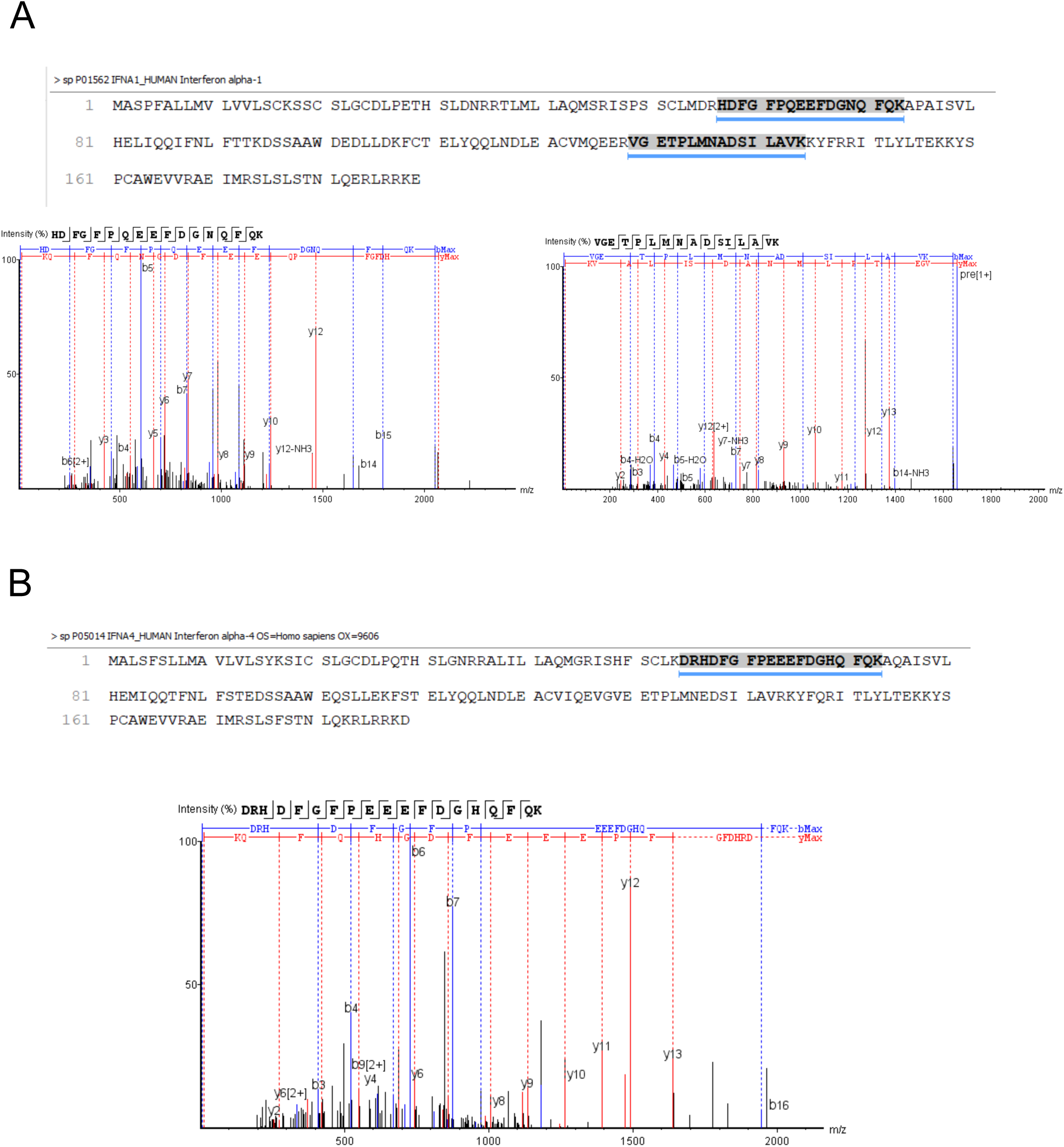
Mass spectrometry profile of protein fraction purified from conditioned media of *IFNAR*s KO Cells. **(A)** Two unique peptides were identified as belonging to IFNα1. **(B)** One unique peptide was identified as belonging to IFNα4.

**Figure S7.**
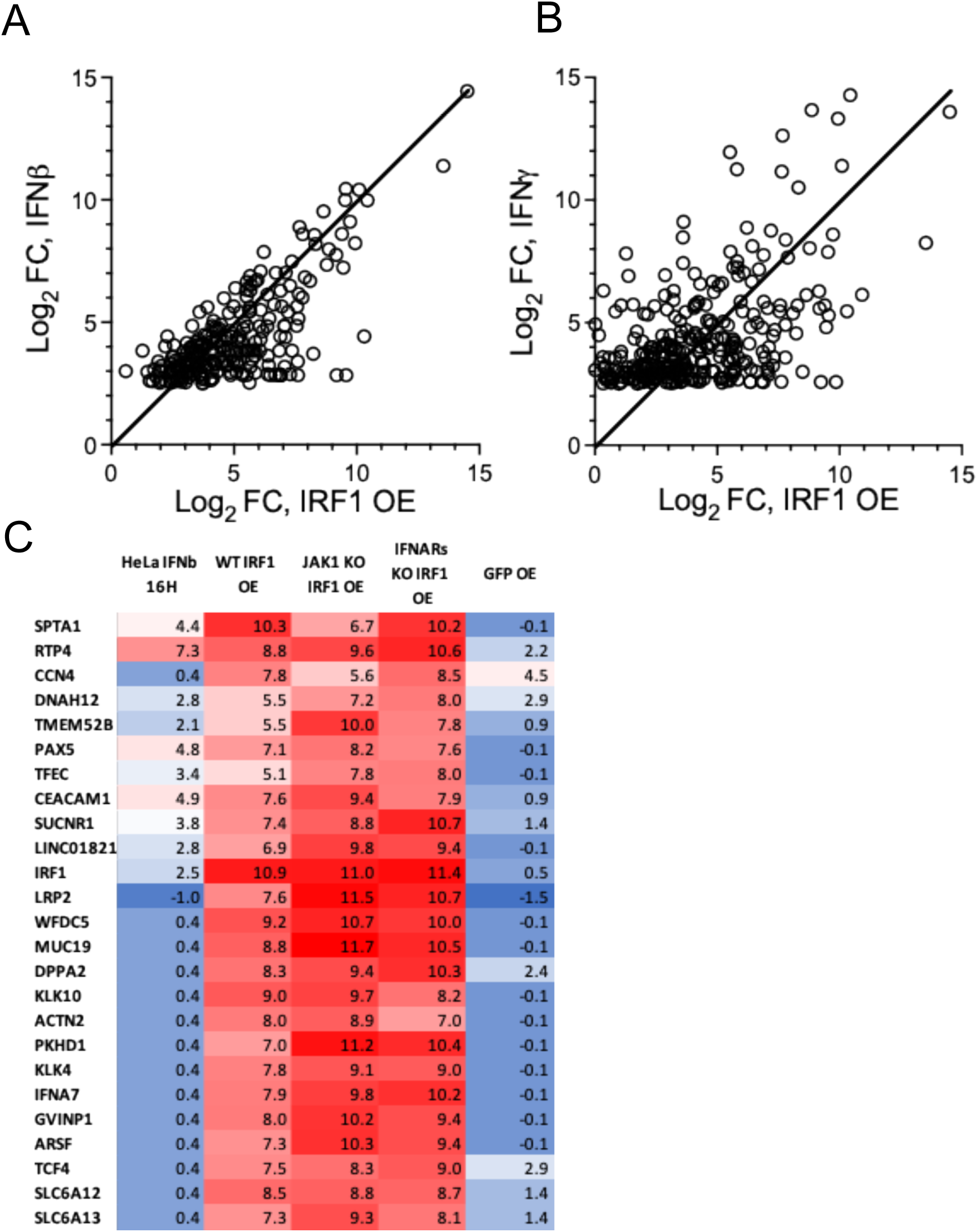
Gene abundance in IRF1 OE cells. **(A, B)** Genes with FC (relative to WT) of >Log_2_ 2.5 upon treatment with 2 nM IFN-β for 16 h **(A)** or 100 nM IFN-γ for 16 h **(B)** plotted versus FC upon IRF1 OE. **(C)** Heatmap of genes extracted from IRF1 OE in WT HeLa cells taken from cluster 3, Fig. 5A under the following conditions: WT IRF1 OE, *JAK1* KO IRF1 OE, and *IFNAR* KO IRF1 OE, as well as WT cells treated with IFN-β (2 nM) for 16 hrs and WT GFP OE used as a transfection control. The values are log(2) relative to WT non-treated. The value of 0.4 represents cases where both the numerator and the denominator are below threshold as calculated by DESeq2. Color scale: blue – lowest, while 4, red – highest.

**Figure S8.**
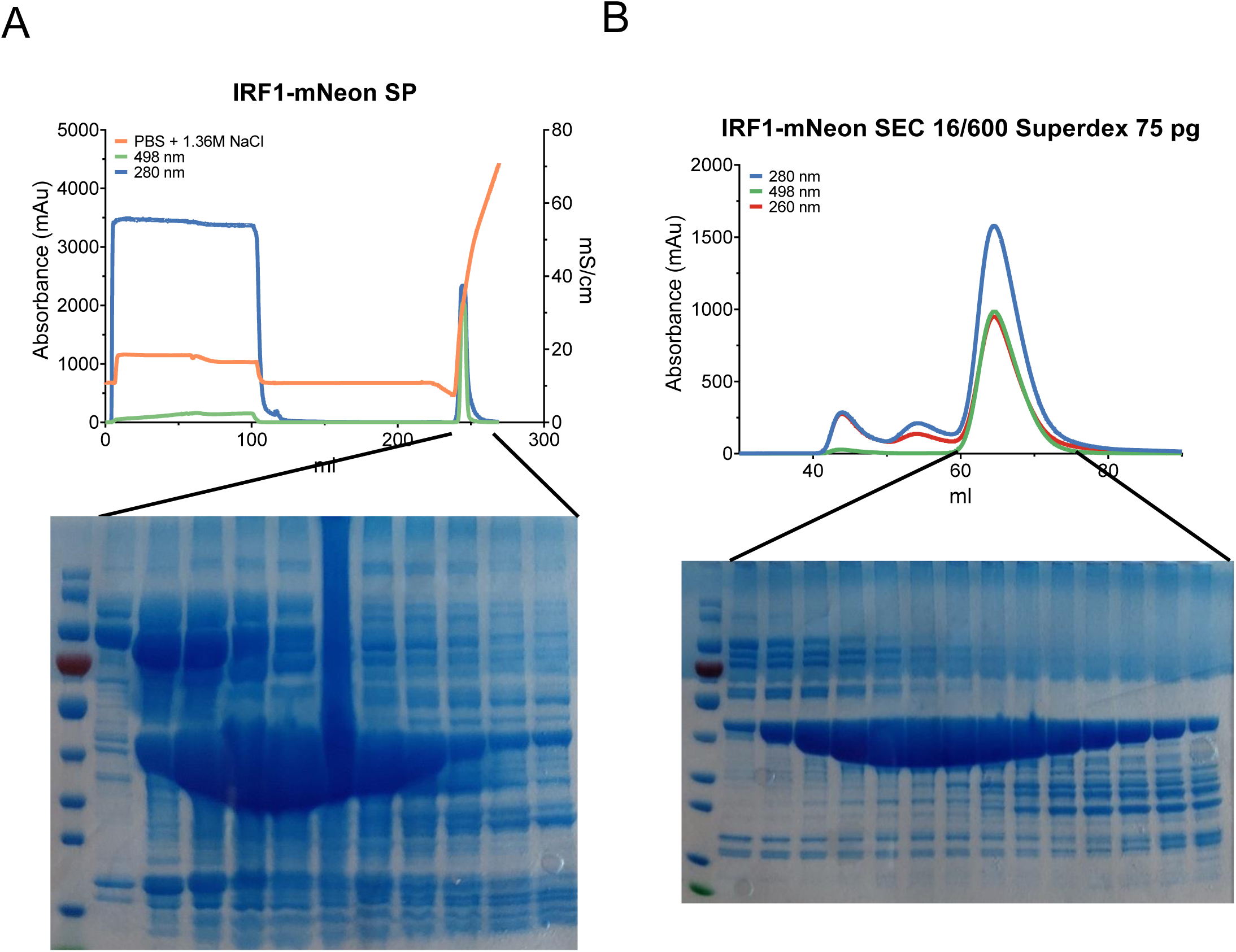
IRF1-mNeon fusion protein purification. **(A)** IRF1-mNeon was purified using HiTrap SP HP cation exchange chromatography. The blue line show absorbance at 280 nm, the green line 498 nm and the orange line show salinity in mS/cm. **(B)** Size exclusion dendrogram of IRF1-mNeon in SEC (superdex 75pg 16/600 chromatography column). The blue line shows absorbance at 280 nm, the red line 260 nm, and the green line 498 nm.

**Figure S9.**
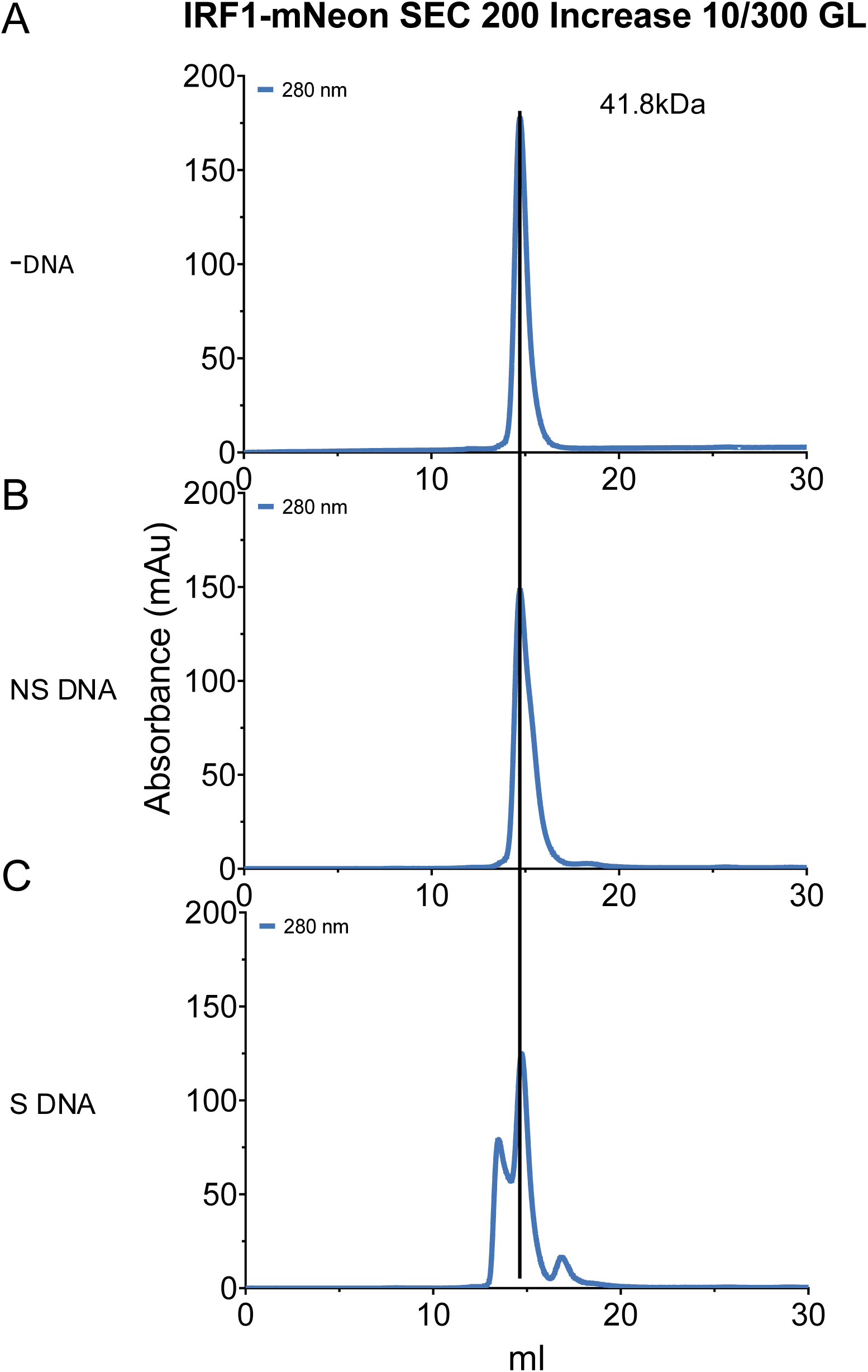
Size exclusion chromatography analysis of IRF1-mNeon complexes. **(A)** –DNA, SEC of IRF1 protein without DNA. **(B)** NS DNA, IRF1 protein incubated for 4 hours with scrambled DNA. **(C)** S DNA, IRF1 protein incubated for 4 hours with a specific DNA. The blue line shows absorbance at 280 nm and the cross-black line show the elution volume of a protein with MW of 41.8 kDa

**Figure S10.**
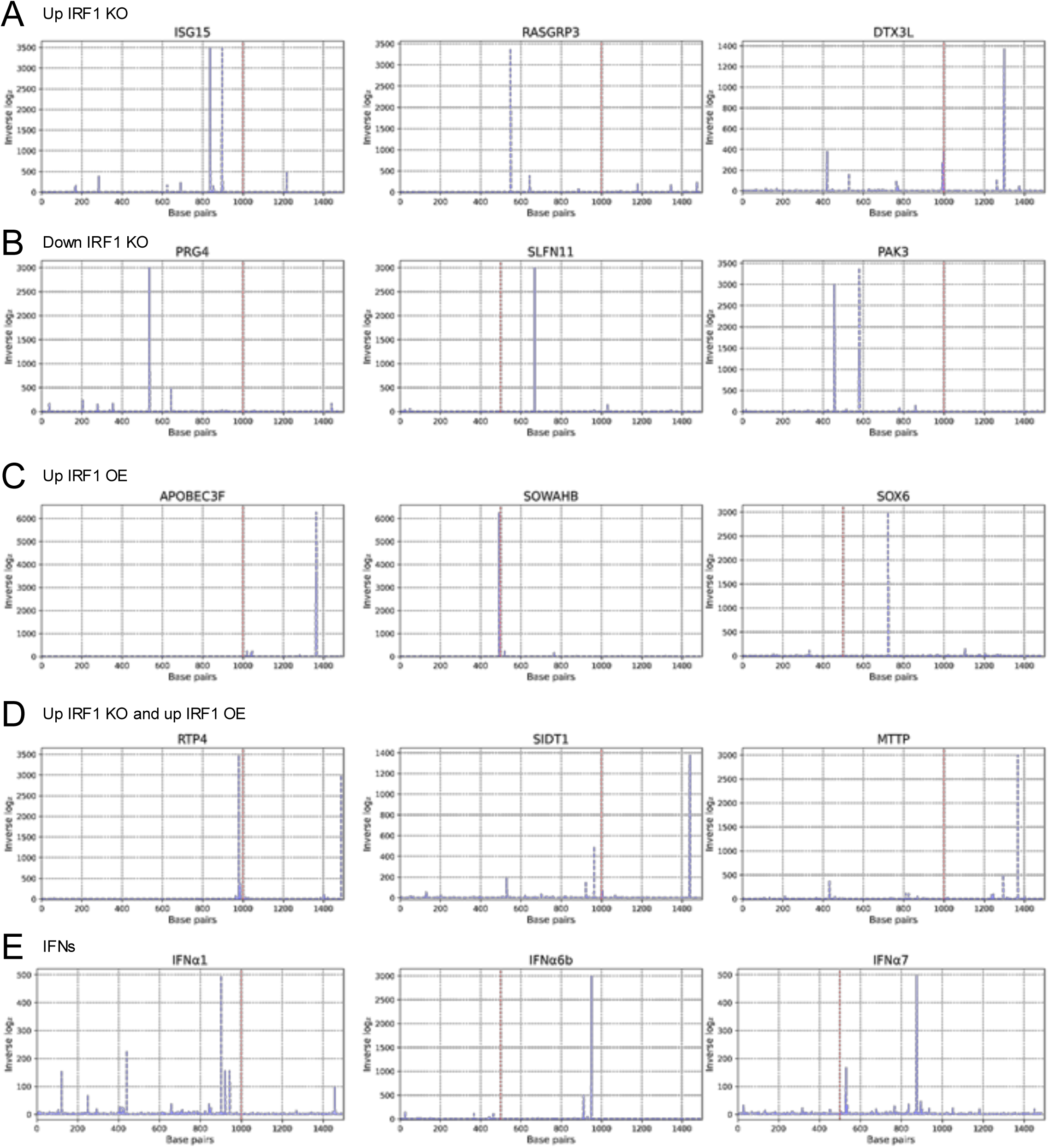
Promoter binding of IRF1 as predicted by our model. (**A-E**) Predicted IRF1 binding peaks across the promoter regions of selected genes that showed significant changes in abundance in the RNA -seq analysis. The promoter regions analyzed are of 1,000 base pairs upstream and 500 base pairs downstream of the transcription start site, marked in red for each gene. Calculated binding affinity is shown as inverse log₂-transformed z-scores (2^z^-score), providing a linear-scale representation of predicted IRF1 DNA interaction strength. (**A**) Increased abundance in the *IRF1* KO. (**B**) Decreased abundance in the *IRF1* KO. (**C**) Increased abundance in IRF1 OE. (**D**) Genes with increased abundance in both *IRF1* KO and IRF1 OE. (**E**) Calculated IRF1 binding sites in promoter regions of type I IFN genes.

**Figure S11.**
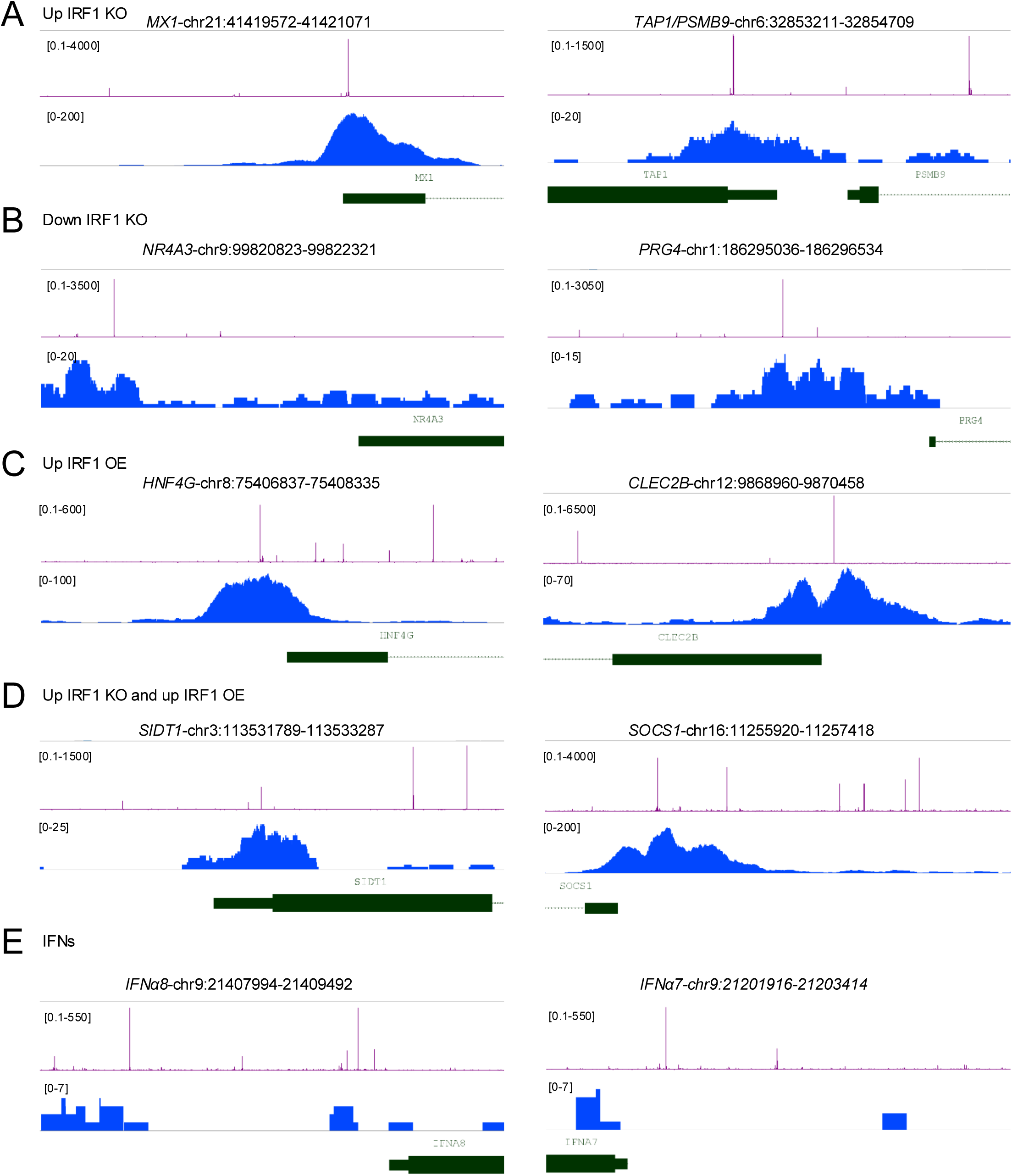
Predicted and observed IRF1 binding in promoter regions of key immune-related genes. (**A-E**) Comparison of predicted IRF1 binding affinity and ChIP-seq signal in promoter regions of genes identified to be controlled by IRF1 according to RNA-seq analysis. The promoter regions span from 1,000 basepairs upstream to 500 basepairs downstream of the transcription start site. Predicted IRF1 binding affinity is shown in purple and was visualized using inverse log₂-transformed z-scores (2^z^-score) to represent the relative strength of predicted interactions on a linear scale. ChIP-seq data, reanalyzed from GEO datasets (GSM6928615, GSM6928616), are represented in blue, showing IRF1 binding coverage across the promoter regions. Gene annotations are highlighted in green.

**Figure S12.**
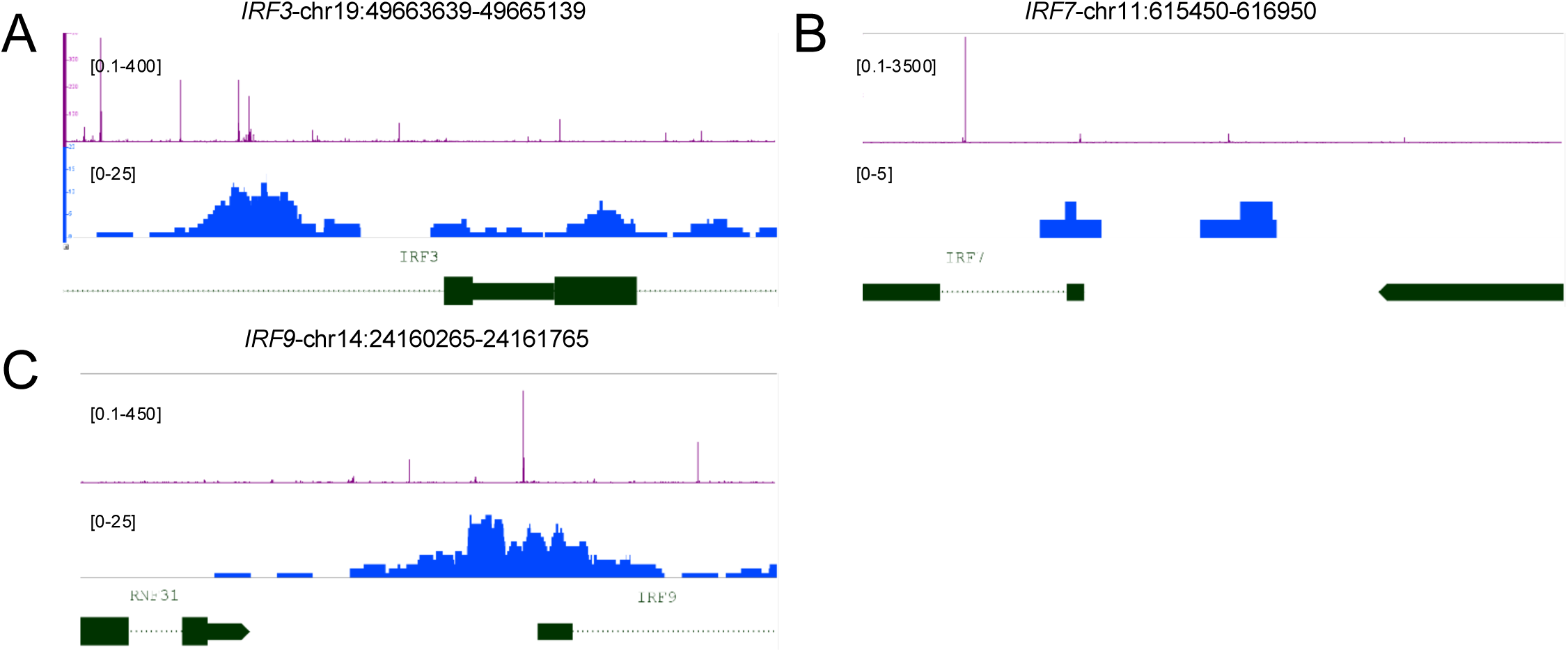
IRF1 binding in promotors of IRF3, IRF7 and IRF9. (A-C) Comparison of predicted IRF1 binding affinity and reanalyzed ChIP-seq coverage in the promoter regions of IRF3 (**A**), IRF7 (**B**), and IRF9 (**C**). Each promoter region spans 1,000 bp upstream and 500 bp downstream of the transcription start site. Predicted IRF1 binding affinity is displayed in purple using inverse log₂-transformed z-scores (2^z^-score), and ChIP-seq coverage is shown in blue. Gene annotations are highlighted in green. These transcription factors were selected based on their altered expression in IRF1 KO cells, as revealed by RNA - seq analysis.

